# Multi-modal and multi-model interrogation of large-scale functional brain networks

**DOI:** 10.1101/2022.12.19.520967

**Authors:** Francesca Castaldo, Francisco Páscoa dos Santos, Ryan C Timms, Joana Cabral, Jakub Vohryzek, Gustavo Deco, Mark Woolrich, Karl Friston, Paul Verschure, Vladimir Litvak

## Abstract

Current whole-brain models are generally tailored to the modelling of a particular modality of data (e.g., fMRI or MEG/EEG). Although different imaging modalities reflect different aspects of neural activity, we hypothesise that this activity arises from common network dynamics. Building on the universal principles of self-organising delay-coupled nonlinear systems, we aim to link distinct electromagnetic and metabolic features of brain activity to the dynamics on the brain’s macroscopic structural connectome.

To jointly predict dynamical and functional connectivity features of distinct signal modalities, we consider two large-scale models generating local short-lived 40 Hz oscillations with various degrees of realism - namely Stuart Landau (SL) and Wilson and Cowan (WC) models. To this end, we measure features of functional connectivity and metastable oscillatory modes (MOMs) in fMRI and MEG signals - and compare them against simulated data.

We show that both models can represent MEG functional connectivity (FC) and functional connectivity dynamics (FCD) to a comparable degree, by varying global coupling and mean conduction time delay. For both models, the omission of delays dramatically decreased the performance. For fMRI, the SL model performed worse for FCD, highlighting the importance of balanced dynamics for the emergence of spatiotemporal patterns of ultra-slow dynamics. Notably, optimal working points varied across modalities and no model was able to achieve a correlation with empirical FC higher than 0.45 across modalities for the same set of parameters. Nonetheless, both displayed the emergence of FC patterns beyond the anatomical framework. Finally, we show that both models can generate MOMs with empirical-like properties.

Our results demonstrate the emergence of static and dynamic properties of neural activity at different timescales from networks of delay-coupled oscillators at 40 Hz. Given the higher dependence of simulated FC on the underlying structural connectivity, we suggest that mesoscale heterogeneities in neural circuitry may be critical for the emergence of parallel cross-modal functional networks and should be accounted for in future modelling endeavours.

## 1 Introduction

The fundamental concepts behind large scale modelling were established as early as 1940s (McCulloch & Pitts, 1943; Shimbel & Rapoport, 1948; Uttley & Matthews, 1955). However, it was only with the advance of technology and neuroimaging techniques — which ensured an unprecedented computational power and spatiotemporal resolution of data — that a procedural framework could be established (Coombes, 2005; Deco et al., 2008; Honey et al., 2007). Large scale modelling aims at finding a balance between complexity and realism, with the goal of explaining data features in a parsimonious and accurate manner. It furnishes a biophysical approach to investigate how the interaction between structural connectivity and intrinsic dynamics gives rise to specific spatiotemporal oscillatory patterns.

To date, inter-regional connectivity patterns and anatomically defined brain regions represent one of the most accurate approximations of the structural organisation — the so called *‘connectome’*: long-range interactions between distant neuronal ensembles are mediated by long axonal projections that can be registered in-vivo and non-invasively using diffusion MRI (Conturo et al., 1999; Hagmann et al., 2008; Sporns et al., 2005). The need for such comprehensive mapping has motivated many researchers to make the structural description more and more detailed, with the aim of describing brain structure on multiple levels and across different species (Alexander et al., 2007; Matthew F. Glasser et al., 2016; Hagmann et al., 2008; Ranzenberger & Snyder, 2022; Zalesky et al., 2010). This detailed structural framework will serve to delineate the space of possibilities in which nodes and their interaction can be modelled as a network.

One of the main questions – in the study of brain oscillations — is to ascertain the relationship between synchronisation mechanisms and collective behaviour and how they depend on coupling strength. Computational models of dynamical systems - such as the models of coupled phase (Kuramoto, 1975), limit cycle (Stuart Landau (Sreenivasan et al., 1987)), and chaotic (Rössler, 1976) oscillators, have been increasingly employed to study the evolving network dynamics emerging from a structured framework (Cabral et al., 2022; Cabral et al., 2011; Cofré et al., 2020; Deco et al., 2008). In 1975, Kuramoto presented a reduced-order model which characterises the within limit-cycle behaviour of nodes, representing the activity of each oscillator (neuron/neural column/cortical area) in terms of its circular phase (Bick et al., 2020; Kuramoto, 1975; Park & Lefebvre, 2020). Moving beyond the limit cycle, in 1987, Andronov and colleagues inspired the implementation of models which include both phase and amplitude modulation (A. et al., 1987). Among those, the complex Stuart-Landau equation has been used to investigate the appearance of an oscillatory mean field from a noisy interacting unit (Pikovsky et al., 2003; Pikovsky & Rosenblum, 2015). Still governed by the same principles — but motivated by neurobiological realism on a different scale — *neural mass models* (NMMs) have provided useful insights into meso- and macroscopic dynamics of populations of interacting excitatory and inhibitory neurons (Beurle & Matthews, 1956; Wilson & Cowan, 1972). In NMMs, each brain region is no longer modelled as a single oscillator (with its own intrinsic frequency), but as a population of neurons whose interaction explains oscillatory dynamics. In both cases, by tuning the model parameters, the system dynamics undergo a phase transition from a noisy to an ordered state. By coupling an ensemble of oscillators/NMMs, the node dynamics can be modelled by the local node/population activity plus the interaction with other regions and noisy fluctuations (Breakspear, 2017).

Extensive research has shown that cortical networks maintain a balance between excitatory and inhibitory activity (Dehghani et al., 2016; Froemke et al., 2007; Sprekeler, 2017; Tao & Poo, 2005; Xue et al., 2014), proven to be beneficial for the cortical function (Litwin-Kumar & Doiron, 2014; Mariño et al., 2005; Páscoa Dos Santos & Verschure, 2021; Rubin et al., 2017; van Vreeswijk & Sompolinsky, 1996; Vogels et al., 2011; Wehr & Zador, 2003). Importantly, such balance is maintained through homeostatic plasticity mechanisms that scale the strength of synapses onto pyramidal neurons to maintain firing rates stable (Ma et al., 2019; Turrigiano, 2011; Turrigiano et al., 1998; Vogels et al., 2011). Furthermore, over the last two decades, modelling studies have highlighted the relevance of incorporating excitatory-inhibitory balance (E/I) to understand the mechanisms that underwrite this balance. One hypothesis is that synaptic plasticity at inhibitory synapses (ISP) plays a key role in balancing E/I inputs and contributes to stabilising the firing rates every time the E/I balance is disrupted by perturbations at the level of incoming excitation. In the context of connectome-based models, the implementation of ISP prevents certain populations of neurons from dominating network behaviour and dynamically adapts synaptic weights to regulate the E/I balance (Landau et al., 2016; Litwin-Kumar & Doiron, 2012). The importance of ISP has been validated on both models of MEG (Abeysuriya et al., 2018) and fMRI (Hellyer et al., 2016), showing its ability in regulating local activity on different timescales and generating more realistic functional patterns.

Large-scale models have shown that they can not only generate an accurate representation of empirical data but also to elucidate structural-functional, and subsequent static-dynamic, relationships. Both structural and functional neuroimaging allow us to model statistical or physical connections in the brain. While structural connectivity refers to the anatomical framework by means of tracing fibre tracts, functional connectivity (FC) is defined as the statistical interaction between disparate brain regions (Friston, 1994). The non-trivial relationship between SC and FC has been increasingly addressed with different modelling approaches, from biophysical (Breakspear, 2017; Deco et al., 2009; Deco et al., 2014; Honey et al., 2007; Pinotsis et al., 2012; Sanz Leon et al., 2013) to statistical, which focus on low-dimensional network diffusion processes or random walks (i.e., (Raj et al., 2020), see (Raj et al., 2022) for a comprehensive review). They share the notion of high-order neural phenomena going beyond the local geometrical clustering but also illustrate how the interplay between local dynamics and the large-scale anatomical framework gives rise to resting-state brain activity (Cabral et al., 2011; Deco, Jirsa, et al., 2013; Deco et al., 2014).

As a proof of concept, strong functional interactions can also exist in absence of structural connections (Hermundstad et al., 2014; Honey et al., 2010) and their spatiotemporal correlation is transiently and dynamically organised (Friston, 1997; Hutchison et al., 2013). Despite these discrepancies, FC is undoubtedly constrained by the anatomical framework. Indeed, on slow time scales FC has been found to indirectly reflect the underlying SC (Honey et al., 2010). In the context of modelling, the FC strongly resembles SC when the system is close to a phase transition and the agreement is best approximated near a bifurcation. This suggests that the optimal working point, linking function to structure, is at the edge of criticality (Cocchi et al., 2017). Although this type of investigation has long been established in the context of fMRI, it has only recently been applied to the field of electromagnetic data (MEG/EEG) (Cabral et al., 2022; Deco, Cabral, et al., 2017; Roberts et al., 2019).

In addition, the dynamic aspect of functional connectivity is becoming increasingly important as the correlation structure shows a rich and dynamic reconfiguration over time (Deco, Kringelbach, et al., 2017; Hutchison et al., 2013) and its alteration reflects cognitive or neurological dysfunction (Bonkhoff et al., 2021; Filippi et al., 2019). Today, large-scale models exist that can explain the possible mechanisms behind the transient motifs of metabolic signals (G. Deco et al., 2021; Deco, Kringelbach, et al., 2017; Vohryzek et al., 2020), but only a few attempts have been made in the context of electrophysiological data (Cabral et al., 2022; Cabral et al., 2014; Deco, Cabral, et al., 2017). Phenomenological models with dynamics on structure allow the behaviour of different models to be compared against empirical observables (Friston & Dolan, 2010). Although different models have attempted to elucidate the mechanisms underlying each of the modalities (see (Glomb et al., 2022) for EEG, (Abeysuriya et al., 2018; Hadida et al., 2018; Raj et al., 2020; Tewarie et al., 2019) for MEG, (Cabral et al., 2022; Cabral et al., 2011; G. Deco et al., 2021; Honey et al., 2007; Roberts et al., 2019; Vohryzek et al., 2020) for fMRI, to date, no large-scale modelling approach has attempted to characterise features across modalities.

In this work, we apply a multi-modal and multi-model approach to recover the underlying neurodynamical genesis of neuroimaging signals. We aim to contribute to the broad repertoire of generative models by proposing a comparative analysis between two large-scale models, identifying advantages and limitations, and testing their applicability in disclosing the network properties of haemodynamic and electrophysiological brain activity. This paper starts with a brief review of the theoretical background for generative modelling of this sort; followed by a description of complementary modelling procedures applied to empirical data. These analyses provide the basis for a comparative evaluation of different modelling strategies and enable us to identify their key functional forms.

## 2 Methods

### 2.1 Phase-amplitude model: Stuart-Landau

The Stuart-Landau (SL) equation (Equation 1) is the canonical form for describing the behaviour of a nonlinear oscillating system near an Andronov-Hopf bifurcation (A. et al., 1987; Cocchi et al., 2017). It describes systems that have a static fixed point but respond to perturbation (i.e., noise, impulse, specific waveform) with an oscillation, which may be damped or self-sustained depending on the operating point of the system with respect to the bifurcation (Supplementary Material (SM), Section I, Figure S1).

Our analysis is based on a system of N=78 SL oscillators coupled in the connectome, considering both the connectivity strength, *C*_*np*_, and the conduction delays, τ_*np*_, between each pair of brain areas *n* and *p*. The conduction delays are defined in proportion to the fibre lengths between brain areas, assuming a homogenous conduction speed *v*, such that τ_*np*_ = *D*_*np*_/*v*, where *D*_*np*_ is the real fibre length detected between brain areas *n* and *p*. To simulate how the activity in node *n* is affected by the behaviour of all other nodes *p* (*p* ∈ N ∧ *p* ≠ *n*), we describe the interaction between nodes in the form:

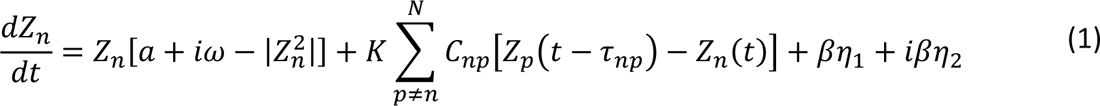

where the complex variable Z_*n*_(t) describes the state of the *n*^tℎ^ oscillator at time t.

The first term in Equation 1 describes the intrinsic dynamics of each unit that is the natural excitability of neuronal assemblies, where ω = 2π ∗ f_f_ is the angular frequency, with f_f_ as the fundamental frequency. As in (Cabral et al., 2022), we set all nodes with identical natural frequency ω_0_ = 2π ∗ 40HZ, representing the ability of a neural mass to engage in gamma-frequency oscillations.

The parameter *a* determines the position of each unit with respect to the limit cycle. For *a* > 0, a stable limit cycle appears via a superciritical Hopf bifurcation, while when *a* < 0 there is only a stable fixed point at the origin Z_*n*_ = 0, so the bifurcation point is at *a* = 0. Importantly, if *a* is negative but sufficiently close to the bifurcation, the system is still weakly attracted to the limit cycle and damped oscillations emerge in response to external input, with a decay time scaled by *a*.

The second term represents the total input received from other brain areas, scaled by parameter *K*, which sets the strength of all network interactions with respect to the intrinsic node dynamics. Because we focus on the nonlinear phenomena introduced by time delays, we model the node-to-node interactions using a particular *linear diffusive coupling*, as the simplest approximation of the general coupling function, considering *delayed* interactions. The last term of Equation 1 represents the real and imaginary part of uncorrelated white noise, where η_1_ and η_2_ are independently drawn from a Gaussian distribution with mean zero and standard deviation β = 0.001. For a detailed exploration and dynamical analysis of SL model see (Cabral et al., 2022; Choe et al., 2010; Powanwe & Longtin, 2021).

### 2.2 Neural mass model

Neural mass-models are mean-field approaches that function under the assumption that the activity of a discrete population of neurons, or neural mass, can be abstracted to its mean, or any other statistic of interest, at a given time (Breakspear, 2017). In our work, to simulate activity of parcellated cortical regions, we make use of one of such approaches: the Wilson-Cowan model of coupled excitatory and inhibitory populations (Wilson & Cowan, 1972). The Wilson-Cowan model describes the firing-rate dynamics of two recurrently connected populations of excitatory (*r*^*E*^) and inhibitory (*r*^*I*^) neurons, being, for this reason, ideal to represent local excitatory-inhibitory balance (Abeysuriya et al., 2018). The dynamics of these two variables can then be described as:

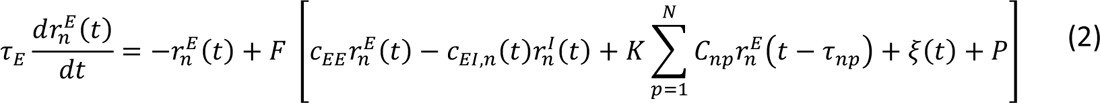

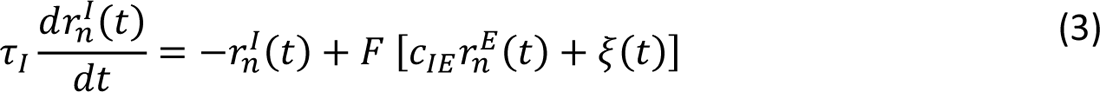

where τ_*E*_ and τ_*I*_ represent the characteristic time constants of the excitatory and inhibitory populations, respectively, *C*_*XY*_ describes the coupling from population y to x (e.g., *C*_*EI*_ represents the inhibitory to excitatory coupling) and *K* is a scaling factor for structural connectivity, hereby referred to as global coupling. *C*_*np*_ represents the structural connection (through white-matter tracts) between nodes *n* and *p* and is based in human structural connectivity data derived from diffusion tensor imaging (see *Structural Connectivity*, Methods). In turn, τ_*np*_, describes the conduction delay between nodes *n* and *p* and is calculated by dividing empirically derived tract lengths by a given conduction speed. Notably, these long-range connections are only implemented between local excitatory populations, in accordance with the evidence that long-range connections in the human cortex are mostly excitatory (Tremblay et al., 2016) and in line with the state-of-the-art in large-scale modelling (Abeysuriya et al., 2018). As in Abeysuriya, Hadida et al., we add a parameter *P* to the description of *r*^*E*^, regulating the excitability of excitatory populations (SM, Section I, Figure S1).

To describe the response of neural masses to external input, we use the function *F*(*x*). Shortly, *F*(*x*) can be roughly equated to the F-I curve of a given population of neurons, and is described as:

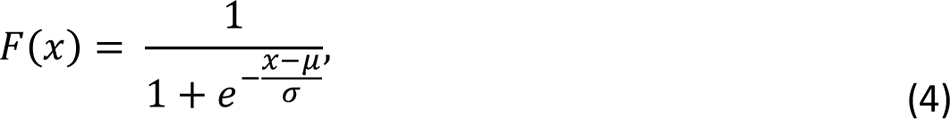

where μ represents the input level at which the neural mass reaches half of its maximum response and can be understood as regulating its excitability, and σ is the approximate slope of the function at that point, being equated to the sensitivity of the neural mass to external input. In addition, both excitatory and inhibitory populations receive uncorrelated additive noise, drawn at each time point from a Gaussian distribution with mean 0 and standard deviation 0.01. For the chosen parameters describing local interactions (*C*_*XY*_) ((Abeysuriya et al., 2018), Table 1), the uncoupled Wilson-Cowan node behaves as a Hopf-Bifurcation between a low-activity steady-state and a limit-cycle (Wilson & Cowan, 1972). Therefore, if the system is close to the bifurcation point, it will transiently exhibit noise-driven oscillations. While the bifurcation point is determined by 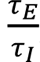, the intrinsic frequency of oscillation depends, instead, on τ_*E*_ τ_*I*_. Since cortical networks are thought to generate intrinsic gamma oscillations through the recurrent interaction between pyramidal cells and fast-spiking inhibitory interneurons (Buzsáki, 2006), we chose τ_*E*_ and τ_*I*_ so that the characteristic frequency of isolated neural masses is within the gamma range (∼40 Hz) (see SM, Section I, Figure S2). In addition, to control the level of input necessary for the phase transition between stable activity and the limit cycle to occur, we regulate the excitability of the neural masses through the parameters μ and *P*. Here, we chose parameters so that an isolated neural mass, with no external input, is in the subcritical regime but sufficiently close to the critical bifurcation point, so that damped oscillations emerge when receiving input from other nodes.

**Table 1.** Table of parameters, values and descriptions for the Wilson-Cowan and Stuart Landau model.

| Wilson-Cowan |  |  | Stuart-Landau |  |  |
| --- | --- | --- | --- | --- | --- |
| Parameter | Value | Description | Parameter | Value | Description |
| $K$ | [0.1, 14] | Global coupling, scaling factor of structural connectivity | $K$ | [4.0, 2000] | Global coupling, scaling factor of structural connectivity |
| Mean Delay | [0, 15] | Mean conduction delay across non-zero connections | Mean Delay | [0, 15] | (ms) Mean conduction delay across non-zero connections |
| $\tau_E$ | 2.5 | (ms) Time constant of excitatory population | $a$ | -5 | Bifurcation parameter |
| $\tau_I$ | 5 | (ms) Time constant of inhibitory population | $\omega$ | $2\pi*40$ | (radians) Intrinsic frequency of oscillation |
| $c_{EE}$ | 3.5 | Recurrent coupling of excitatory populations | $\beta$ | 0.001 | Standard deviation of additive gaussian noise |
| $c_{IE}$ | 3.75 | Coupling from excitatory to inhibitory populations | | | |
| $P$ | 0.31 | Adjusts excitability of excitatory population | | | |
| $\mu$ | 1 | Firing threshold of activation function $F(x)$ | | | |
| $\sigma$ | 0.25 | Sensitivity of activation function $F(x)$ | | | |
| $\tau_{homeo}$ | 2500 | (ms) Time constant of homeostatic plasticity | | | |
| $\rho$ | 0.22 | target firing rate of homeostatic plasticity | | | |
| $\xi$ | N<br>(0,0.01) | Additive gaussian noise | | | |

### 2.3 Homeostatic plasticity

To study the effect of balancing excitation and inhibition at the level of single Wilson-Cowan nodes, we implemented a homeostatic mechanism known as synaptic scaling of inhibitory synapses (Maffei & Turrigiano, 2008; Vogels et al., 2011). This type of approach has been previously implemented in large-scale models of the human cortex (Abeysuriya et al., 2018) and inhibitory synaptic scaling has been shown to play an essential role in cortical function and homeostasis (Ma et al., 2019). Therefore, we implemented homeostatic plasticity to adjust local inhibitory weights so that excitatory activity (*r*^*E*^) is corrected towards a given target firing rate (ρ). Therefore, the dynamics of local inhibitory couplings *C*_*EI,i*_ can be described by the following equation, following (Vogels et al., 2011):

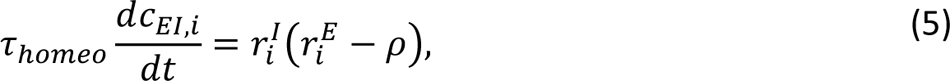

where τ_homeo_ is the time constant of plasticity. In the cortex, the homeostatic mechanisms that are responsible for the maintenance of excitatory-inhibitory balance are known to operate in slow timescales, often hours to days (Turrigiano, 2011). However, to ensure the computational tractability of our simulations, we chose τ_homeo_ = 2.5s. This choice is unlikely to affect our results significantly, since the influence of τ_homeo_ in our system is in determining how quickly local inhibitory weights evolve towards their steady state. In fact, if homeostatic plasticity is sufficiently slow to be decoupled from fast dynamics of intrinsic oscillations, *C*_*EI*_ will reach nearly the same steady state, independently of the time constant (SM, Section I, Figure S3). We also ran simulations not considering homeostatic plasticity to pursue a comparative analysis (see SM, Section I, Figure S4).

### 2.4 Models Parameters

### 2.5 Hemodynamic model

To extract a blood-oxygenation-level-dependent (BOLD) signal equivalent from our simulations, we make use of a forward hemodynamic model (Friston et al., 2000), that incorporates the Balloon-Windkessel model (Friston et al., 2003). In short, hemodynamic models describe how population firing rates (a proxy for neuronal activity) influence the vasculature, which in turn affects blood flow, inducing changes in blood vessel volume and deoxyhemoglobin content, which underlie BOLD signals. In our work, we chose to use the activity from the excitatory populations (*r*^*E*^) only as the input of the Balloon-Windkessel model. This choice is unlikely to influence final results, given the similarity between *r*^*E*^ and *r*^*I*^ in the Wilson-Cowan model (SM, Section I, Figure S1). All of the parameters were taken from (Friston et al., 2003). In addition, we downsample simulated BOLD signals to a period of 0.72s to equate the sampling frequency of the empirical data used in this work (see *fMRI*, Methods).

### 2.6 Model optimisation

We performed model optimization by treating the global coupling (K) and mean delay (mean tract length divided by conduction velocity) as free parameters for both models. For the Wilson-Cowan model, we fixed the target firing rate (ρ) of homeostatic plasticity at 0.22. We also ran simulations for different values of ρ (see SM, Section I, Figure S5). For both models, we performed a grid search over the mentioned free parameters, with 25 logarithmically spaced values of K and 16 values of mean delays in steps of 1ms. Parameter ranges can be consulted in Table 1. For the SL model, simulations with the two highest values of K explored led to instability and results are, therefore, not presented.

For the WC simulations, due to the dynamics of homeostatic plasticity, there was a need to ensure that local inhibitory weights reached a stable or quasi-stable steady state before activity was recorded. Therefore, during simulations, we record *C*_*EI*_ weights every 10s, enough to capture their slow dynamics. We then monitor the evolution of *C*_*EI*_ and allow simulations to run for either 500 minutes of simulation time or until local weights converged to a steady state for all network nodes, evaluated via the condition described in supplementary material (SM, Section I, Figure S6). After ensuring that *C*_*EI*_ reached a steady state, we disable plasticity and record 20 minutes of model activity. Although the slow dynamics of E-I homeostasis prevent it from interacting with the fast dynamics of neural activity, we follow this procedure similarly to previous approaches (Abeysuriya et al., 2018; Hellyer et al., 2016). Regarding the SL model, we run and record 20 minutes of simulation. We ran simulations with an integration time step of 0.2ms.

For both models, after obtaining 20 minutes of simulations, we passed the simulated activity through a haemodynamic model to obtain a synthetic BOLD signal and remove the first and last 2.5s to avoid boundary effects, thus obtaining 15 minutes of BOLD signal timeseries (Friston et al., 2000). We then stored the last 50 seconds of simulated activity (downsampled to 250 Hz) as MEG signals. We compared simulated and empirical FC matrices (see *Data and Model Analysis*, Methods) through the correlation coefficient between their upper triangular parts, and FCD distributions, through the Kolmogorov-Smirnov (KS) distance between them (Lopes et al., 2007). To identify an optimal working point for each model and each measured modality (BOLD, MEG theta, MEG alpha and MEG beta – see below for details), we iterate over a range of thresholds for FC correlation (*CC* ≥ tℎ_*FC*_) and FCD KS distance (*KS* ≤ tℎ_*FCD*_)) and identify the maximum value of tℎ_*FC*_ − tℎ_*FCD*_ for which both conditions can be satisfied by at least one point in the parameter space (see SM, Section II, Figure S7). We then define our model’s working point, for each modality, as the combination of parameters that satisfies both thresholds. Since we primarily focus on the representation of relevant FC patterns, we impose 0.45 as the minimum tℎ_*FC*_.

### 2.7 Data collection and processing

#### 2.7.1 Ethics statement

All human data used in this study is from the public repository of the Human connectome Project (HCP) (https://www.humanconnectome.org), which is distributed in compliance with international ethical guidelines.

#### 2.7.2 Structural Connectivity

The *NxN* matrices of structural connectivity, C, and distances, D, used in the brain network model were computed from diffusion spectrum and T2-weighted Magnetic Resonance Imaging (MRI) data obtained from 32 healthy participants scanned at the Massachusetts General Hospital centre for the Human connectome Project (http://www.humanconnectome.org/).

Briefly, the data were processed using a generalised q-sampling imaging algorithm implemented in DSI Studio (http://dsi-studio.labsolver.org). A white-matter mask, derived from the segmentation of the T2-weighted anatomical images, was used to co-register the images to the b0 image of the diffusion data using the SPM12 toolbox (https://www.fil.ion.ucl.ac.uk/spm/software/spm12/). In each participant, 200,000 fibres were sampled within the white-matter mask. Fibres were transformed into Montreal Neurological Institute (MNI) space using Lead-DBS (Horn & Blankenburg, 2016).

The connectivity matrix C was obtained by counting the number of fibres detected between each pair of *N=78* brain areas defined in the Automated Anatomical Labelling (AAL) parcellation scheme. Similarly, the distance matrix D was obtained by computing the mean length of all fibres detected between each pair of *N=78* cortical brain areas.

#### 2.7.3 fMRI

Empirical fMRI data from healthy subjects was obtained from the public database of the Human Connectome Project (HCP), WU-Minn Consortium (Principal Investigators: David Van Essen and Kamil Ugurbil; 1U54MH091657) funded by the 16 NIH Institutes and Centers that support the NIH Blueprint for Neuroscience Research, and by the McDonnell Center for Systems Neuroscience at Washington University(Van Essen et al., 2013). More specifically, this data was obtained from 99 unrelated subjects (mean age 29.5, 55% females). Each subject underwent four resting-state fMRI sessions of around 14.5 minutes on a 3-T connectome Skyra scanner (Siemens) with the following parameters: TR = 0.72 s, echo time = 33.1 ms, field of view = 208×180mm, flip angle = 52°, multiband factor = 8, echo time = 33.1 with 2×2×2 isotropic voxels with 72 slices and alternated LR/RL phase encoding. For further details, on the standard processing pipeline for HCP data, please consult (M. F. Glasser et al., 2016) and https://www.humanconnectome.org/study/hcp-young-adult/data-releases. In this work, we use the data from the first session of the first day of scanning only.

We further parcellate voxel-based data into 90 anatomically segregated cortical and subcortical regions, excluding the cerebellum, using the Anatomic Automatic Labeling (AAL) atlas. Given that we focus on cortical dynamics, we exclude the 12 subcortical regions, and perform a voxel-wise average of BOLD signals associated with each of the remaining 78 cortical regions, reducing the size of our data to 78 areas x 1200 TR timeseries.

#### 2.7.4 MEG

Pre-processed sensor level MEG data, along with a defaced structural MRI and the appropriate affine transformation matrix mapping between the MRI and MEG spaces were downloaded from the HCP data repository (Wu-Minn HCP 1200 Subjects Data Release). Each of the 89 subjects underwent 6-minute resting state scans (where they were instructed to lie still and keep their eyes open), giving a total of 267 datasets. Full details of the pre-processing steps performed by the HCP team can be found in the HCP manual (https://www.humanconnectome.org/storage/app/media/documentation/s1200/HCP_S1200_Release_Reference_Manual.pdf).

All processing steps were carried out in FieldTrip (Oostenveld et al., 2011) in MATLAB 2021b. The anatomical MRI was linearly transformed from the native MRI space to the MEG scanner space, before being segmented into grey matter, white matter, and cerebral spinal fluid. This segmentation informed the construction of a Nolte single shell head model (Nolte, 2003). A common template array of voxels (isotopically distributed on a grid with 8mm separation, confined to lie within the brain) was non-linearly aligned from MNI space to each of the individual subject’s anatomical images using SPM8’s “old normalise” function (Ashburner & Friston, 2005). This meant that there was a “standard” source model used in the pipeline, with one-to-one correspondence between sources across subjects.

Nearest-neighbour interpolations between this template grid and the atlases that we used in this study were used, allowing us to parcellate voxels into anatomically defined brain regions. A volumetric lead field matrix was calculated for each of the voxel locations. We collapsed the rank of the lead field for each voxel from three to two by executing a singular value decomposition (SVD), thus eliminating any sensitivity to the weakly contributing radial component of the lead field (Ahlfors et al., 2010; Hämäläinen et al., 1993).

The pre-processed sensor level MEG recordings were further band-pass filtered between 1 and 45Hz and downsampled to 250Hz. These data were used to construct a covariance matrix for the construction of linearly constrained minimum variance (LCMV) beamformer weights (Van Veen et al., 1997). This matrix was regularised by adding 1% of the average eigenvalue to the diagonal to improve numerical stability and boost the reconstruction accuracy of the estimated time series (Van Veen et al., 1997). At each voxel location, a separate SVD was run on the 3-dimensional vector time series to extract the optimal lead field orientation in order to maximise the SNR of beamformer weights (Sekihara et al., 2004), thus collapsing the 3 element timeseries to a single time series for each voxel.

### 2.8 Data and Model Analysis

#### 2.8.1 fMRI FC

To compute functional connectivity (FC) from BOLD signals, both empirical and simulated, we calculate pair-wise correlations between all individual timeseries from each of the 78 cortical areas of the AAL atlas, using the Pearson’s correlation coefficient. We then averaged FC over the 99 subject-specific correlation matrices to obtain a 78×78 empirical FC matrix, against which simulated FC matrices can be compared.

#### 2.8.2 MEG FC

Upon obtaining estimates for the neural source currents, data were parcellated into nodes pertaining to each atlas. The first principal component was extracted from all voxels within each ROI. Data were then corrected for spurious correlations arising from source leakage between brain regions by means of symmetric orthogonalization (Colclough et al., 2015). After bandpass filtering the data, we took the analytical signal of the Hilbert envelope for all brain regions and derived whole brain functional connectivity networks by calculating the pair-wise Pearson correlation between each network node. Note that this was done on both the broadband and frequency-specific bands of activity. Finally, we calculated the average amplitude envelope FC matrix over all subjects and sessions. See SM, section IV, for further details on amplitude envelope correlation, MEG source leakage correction and beamforming methods.

#### 2.8.3 fMRI Functional Connectivity Dynamics

While research has mostly focused of the static properties of FC, recent results show that functional connectivity exhibits complex spatiotemporal dynamics, with the transient reinstatement of connectivity states (Deco, Kringelbach, et al., 2017). Here, to evaluate functional connectivity dynamics (FCD), we make use of the method presented in (Abeysuriya et al., 2018; G. Deco et al., 2021; Deco, Kringelbach, et al., 2017). We first split data in N_T_ windows of 80 samples (∼1 minute) with 80% overlap and compute FC within each window following the method described in the previous section. Then, for all pairs of windows, we compute the Pearson’s correlation between the upper triangle of their respective FC matrices. We thereby obtain an N_T_ x N_T_ matrix containing all pairwise correlations between the windowed FC matrices. We then concatenate the values in FCD matrices across subjects to obtain an empirical distribution, against which we compare FCD distributions from each simulation.

#### 2.8.4 MEG Functional Connectivity Dynamics

To compute FCD from MEG signals we make use of a similar method to that described in the previous section, with minor changes, given the nature of MEG signals. First, we filter MEG data at three frequencies of interest (theta: 4-8 Hz, alpha: 8-13 Hz, beta: 13-30 Hz) and compute frequency-specific amplitude envelopes as described in the analysis section of the methods. Then, we split each time-series into windows of 500 samples (2s), with 50% overlap. While the appropriate window size for the calculation of FCD in MEG signals is not yet clear, partly due to the heterogeneity in timescales of the emergence of spatiotemporal MEG patterns, our chosen value is within the range previously used in literature (Liuzzi et al., 2019). Finally, we follow the procedure described for BOLD signals, for each frequency band. From the empirical MEG data, we then obtain frequency-specific distributions of FCD.

#### 2.8.5 Metastable Oscillatory Modes

Previous results suggest that coupled oscillators with delayed interactions give rise to the emergence of metastable oscillatory modes (MOMs) (Cabral et al., 2022). These MOMs consist in transient moments of synchronization between clusters of nodes in a network at frequencies that are lower than the intrinsic frequency of oscillation of uncoupled nodes.

To detect MOMs in both empirical and simulated fMRI and MEG data we first filter timeseries at the bands of interest (fMRI: 0.008-0.08 Hz, MEG theta: 4-8 Hz, MEG alpha: 8-13 Hz, MEG beta: 13-30 Hz). Then, we calculate the respective Hilbert envelopes by computing the absolute value of the Hilbert transform of timeseries from each individual area. Hilbert envelopes are then Z-scored (Z = (*x* − μ)/σ, where *x* is the Hilbert envelope, μ its mean and σ its standard deviation) and a threshold of 2 is applied for the detection of MOMs. While the threshold is arbitrary, assuming that data is normally distributed, a value of 2 represents the threshold above which an incursion of the signal is distinct from noise with a significance level of p<0.05 (Hellyer et al., 2016). Different thresholds were tested, leading to the same qualitative results when comparing simulated and empirical data (SM, Section II, Figure S11).

While in the original approach MOMs were detected using a threshold derived from activity of models without delayed interactions (Cabral et al., 2022), we chose instead to threshold timeseries against their own standard-deviation. We followed this approach to compare the properties of MOMs from simulated and empirical results, since the original method does not allow for the detection of MOMs in empirical data. Similar methods have been applied to the detection of neural avalanches in MEG and MRI data (Hellyer et al., 2016; Sorrentino et al., 2021).

To quantify the properties of MOMs, similarly to (Cabral et al., 2022), we use of the following metrics:

1. **Size:** number of areas with amplitude higher than threshold at a given point in time
2. **Duration:** continuous time interval during which an amplitude timeseries is higher than threshold
3. **Occupancy:** proportion of time a network exhibits oscillations with above-threshold amplitude.

## 3 Results

In order to investigate the spontaneous dynamics observed in resting state fMRI and MEG data of healthy individuals, we used two generative brain network models with different degree of realism, namely the Stuart Landau model – based on a system of delayed coupled oscillators, and the extended version of Wilson and Cowan model – based on a system of coupled excitatory and inhibitory neural populations including delays and homeostatic inhibitory plasticity. We focus on functional connectivity and spatiotemporal data features and show the ability of large-scale models in reproducing key empirical patterns across modalities. Additionally, by linking endogenous oscillations and neuroanatomic structure, we explore the emergence of itinerant dynamics in both models and how it relates to transient properties of multiresolution brain activity. The pipeline overview is illustrated in Figure 1.

**Figure 1.**
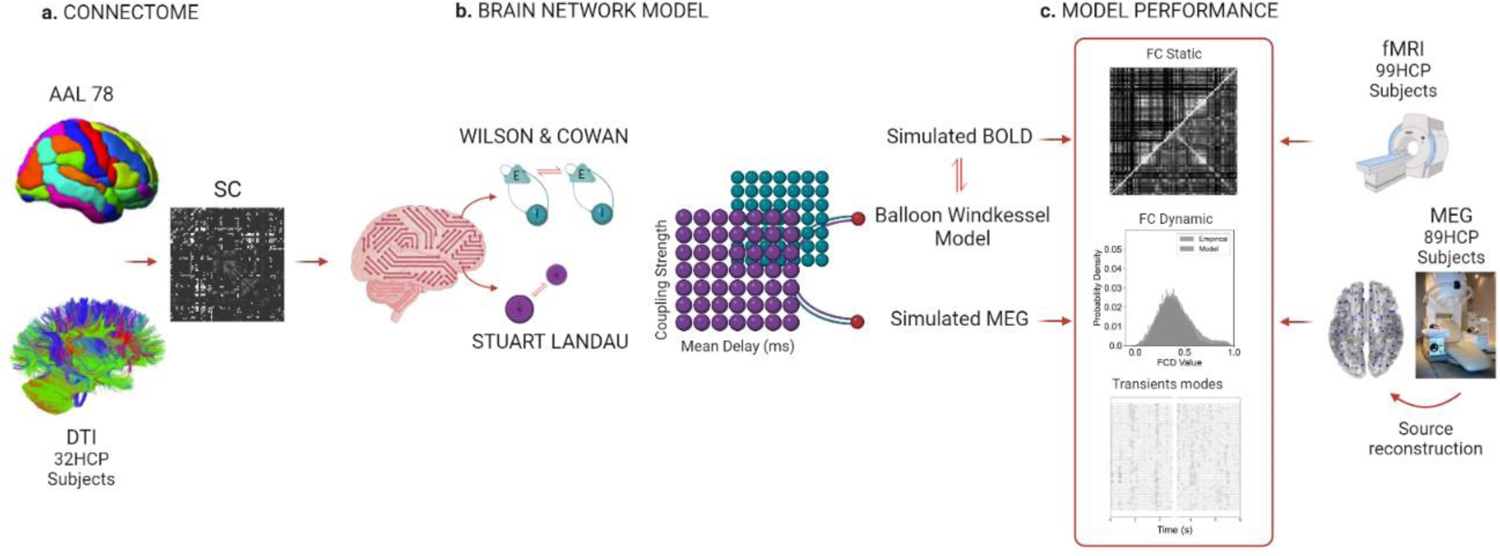
**a.** To build our structural connectivity (SC) we use averaged diffusion tensor imaging (DTI), generated by delineating the white matter fibres orientation of 32 healthy subjects and a cortical parcellation (AAL) for partitioning the grey matter interface into 78 Region of Interests (ROIs). In the final SC graph, each ROI becomes a node and fibres become edges. **b.** We use this connectome to inform both phenomenological models. Both models are characterised by non-linear differential equations, whose parameters are tuned according to physiological plausibility to generate the oscillatory patterns observed empirically. Building on previous findings, the intrinsic frequency of all units is set at ω = 40Hz and each unit is perturbed with uncorrelated white noise for both models (Cabral, Castaldo et al. 2022). In this study, we optimised two global parameters; namely, the coupling strength and the mean conduction delay, which were varied over specific ranges to best explain the empirical data features. For each combination of these two parameters, the models generate an oscillatory pattern. To create a blood-oxygenation-level-dependent (BOLD) signal from our simulations, a forward hemodynamic model is implemented, while we do not apply any additional steps to represent the simulated MEG signals. **c.** Both simulated and empirical signals follow the same pre-processing and analysis steps before being compared: for each combination of global parameters, we compute and compare the models’ and empirical functional connectivity, functional connectivity dynamics and properties of metastable oscillatory modes (see Methods for details).

### 3.1 Suitability of two large-scale models for simultaneously representing empirical motifs of static and dynamic functional connectivity

Both models reasonably approximate patterns of connectivity and temporal variability observed in empirical BOLD fMRI and MEG signals. Yet, delays have a different impact depending on which signal is reproduced. As shown in Figure 2b and Figure 3b, delays play a key role in shaping the frequency content of MEG connectivity profiles and this observation holds for both models.

**Figure 2.**
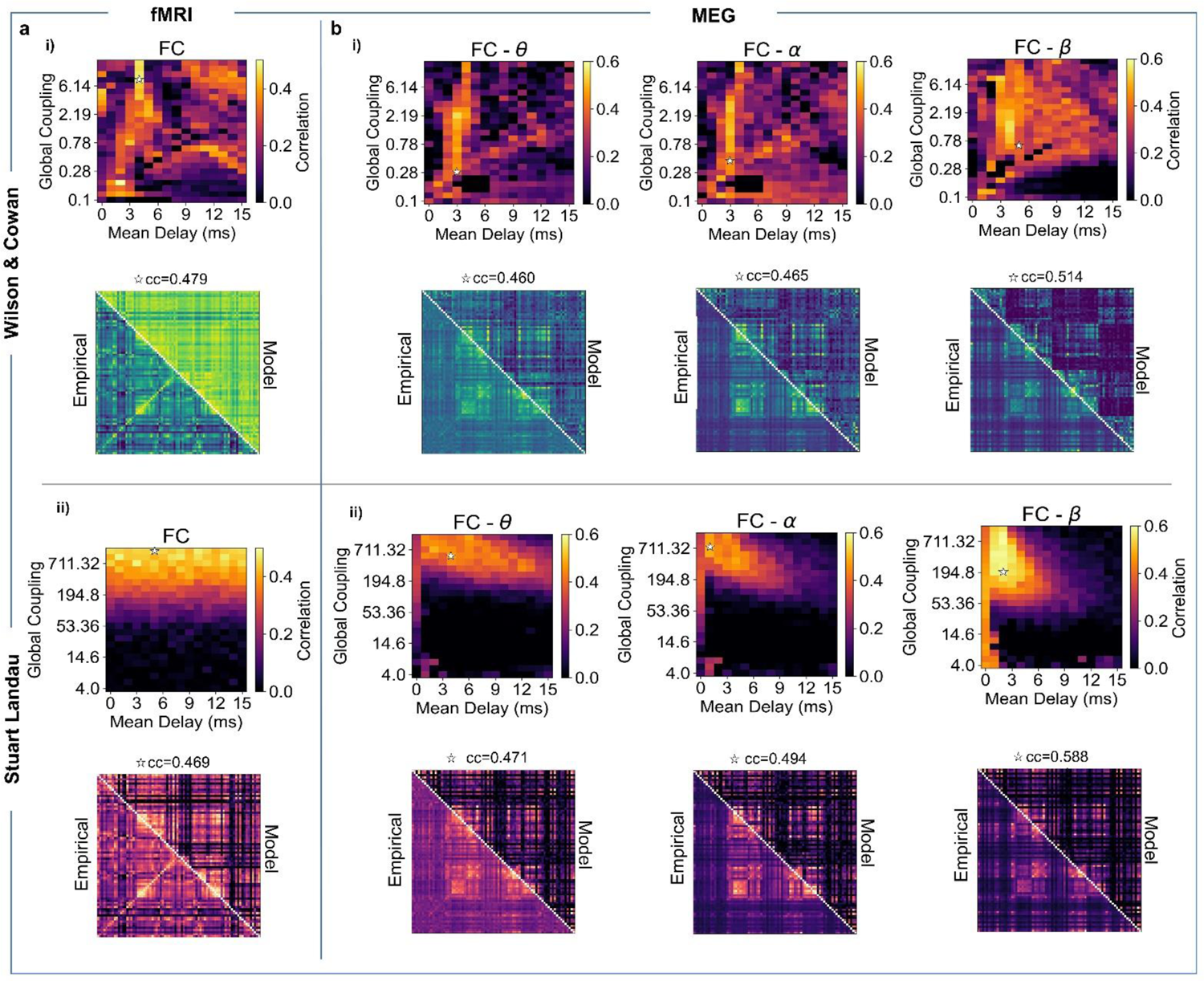
Frequency-specific connectivity patterns emerge from the model network parameter space. **a.** Model performance in explaining empirical BOLD fMRI static connectivity measures: Pearson correlation between BOLD fMRI FC (averaged across 99 HCP participants) and simulated FC for each pair of parameters (Mean Delay and Global Coupling) for WC and SL model. **i)** *Top* - For the WC model, the selected optimal point (white star) for BOLD fMRI is C=7.55, MD=4ms, with correlation of c= 0.479 and ks-distance value of ks=0.107. *Bottom* - Empirical and simulated fMRI BOLD FC for 78 AAL cortical brain areas in the optimal point. **ii)** *Top* - For the SL model, the optimal point for BOLD fMRI is C=1194.16, MD=5ms, with correlation of c= 0.469 and ks-distance value of ks=0.489. *Bottom* - Empirical and simulated fMRI BOLD FC for 78 AAL cortical brain areas in the optimal points. **b.** Model performance in representing empirical MEG connectivity measures: Pearson correlation between Hilbert envelope FC of MEG (averaged across 89 HCP participants) and simulated Hilbert envelope FC for each pair of parameters, for theta [4-8 Hz] (left), alpha [8-13Hz] (middle), beta [13-30Hz] (right) for WC and SL model. **i)** *Top* - The selected WC parameters for MEG are C=0.34, MD=3ms for theta; C=0.42, MD=3ms for alpha; C=0.78, MD=4ms for beta with correlation values of c_theta=0.460, c_alpha=0.465, c_beta=0.514 and ks-distance values of ks_theta=0.084, ks_alpha=0.194, ks_beta=0.395. *Bottom -* Empirical and simulated Hilbert envelope MEG FC matrices for theta (left), alpha (middle), and beta (right) band in the selected optimal point. **ii)** *Top* - The selected SL model parameter combinations for MEG are C=549, MD=4ms for theta; C=711.32, MD=1ms for alpha; C=194.8, MD=2ms for beta with correlation values of c_theta=0.471, c_alpha=0.494, c_beta=0.588 and ks-distance values of ks_theta=0.168, ks_alpha=0.104, ks_beta=0.149. *Bottom -* Empirical and simulated Hilbert envelope MEG FC matrices for theta (left), alpha (middle), and beta (right) band in the optimal point. White stars indicate the model working points, chosen through simultaneous optimization for the representation of empirical FC and FCD, as described in the Methods section.

**Figure 3.**
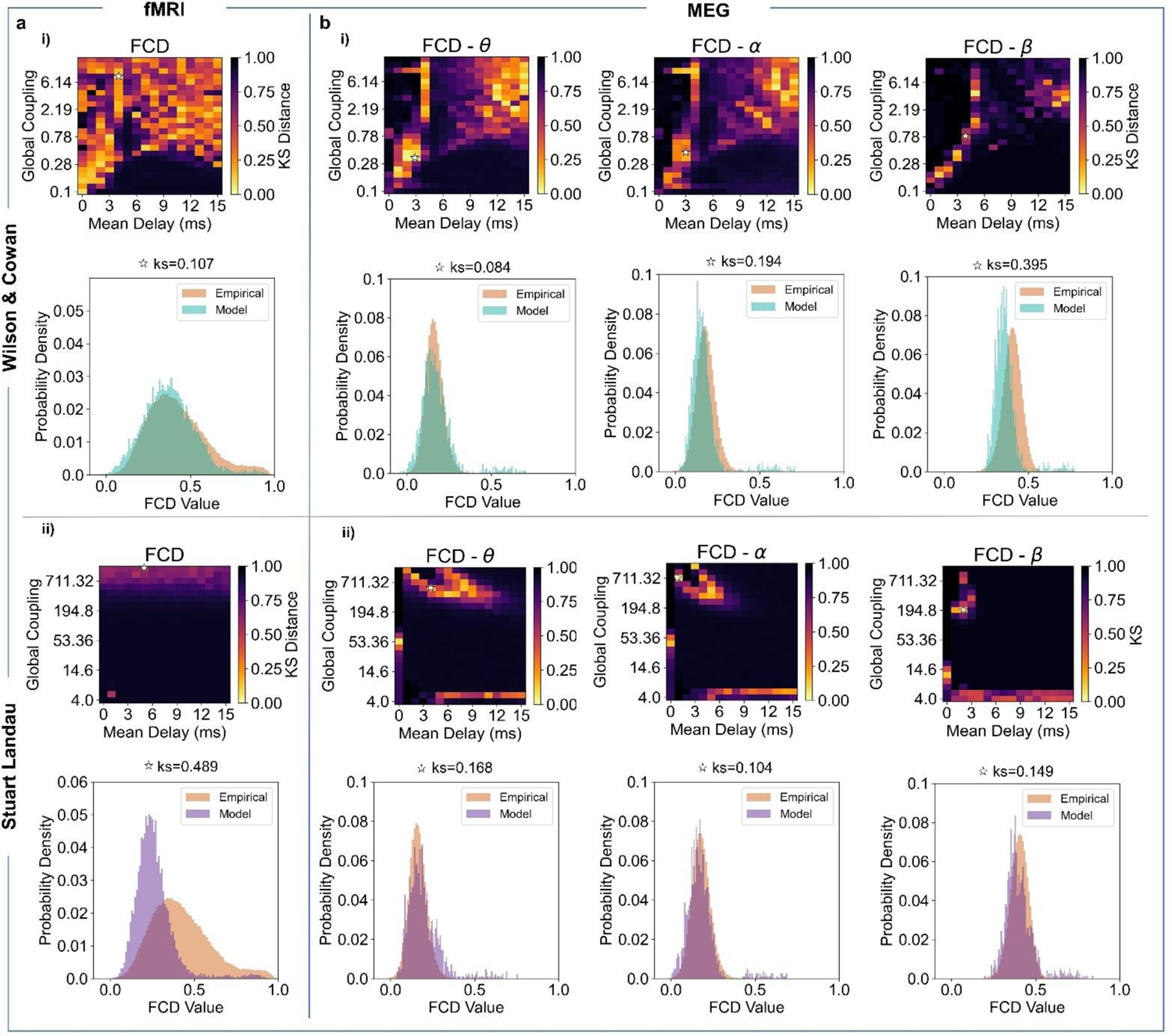
Dynamical spatiotemporal patterns emerge from the model network parameter space. **a.** Model performance in representing empirical BOLD fMRI dynamical connectivity measures: Kolmogorov-Smirnov (KS) distance between empirical BOLD fMRI FCD histograms (averaged across 99 HCP participants) and simulated FCD histograms for each pair of parameters (Mean Delay and Global Coupling) for WC and SL model. **i)** *Top -* For the WC model, the optimal parameters (white star) for BOLD fMRI are C=7.55, MD=4ms, with ks-distance value of ks=0.107. *Bottom -* Empirical and simulated fMRI BOLD FCD distribution for 78 AAL cortical brain areas in the optimal point. **ii)** *Top -* For the SL model, the optimal points for BOLD fMRI are C=1194.16, MD=5ms, with ks-distance value of ks=0.489. *Bottom -* Empirical and simulated fMRI BOLD FCD distribution for 78 AAL cortical brain areas in the optimal point. **b.** Model performance in representing empirical MEG connectivity measures: Kolmogorov-Smirnov (KS) distance between empirical Hilbert envelope MEG FCD histograms (averaged across

However, when it comes to assessing the key patterns seen in the fMRI data, unlike in the WC model, delays do not affect the SL oscillatory dynamics (Figures 2a, 3a). The static functional connectivity and dynamical profiles are plotted for those values in the parameter space that optimise both FC and FCD simultaneously, displayed as white stars in the parameter space (see Methods).

Nevertheless, for the WC model, there is a region of parameter space where the simulated metabolic patterns approximate the empirical patterns, when delays are set to zero (Figures 2-3a, i). This result aligns with the literature on fMRI modelling, which succeeded in approximating haemodynamic activity without considering delays (Deco, Kringelbach, et al., 2017). However, it should be noted that when accounting for local dynamics — whose intrinsic frequency is higher than the intrinsic frequency of low-frequency signals (i.e., fMRI) – delays have been shown to induce much richer and realistic dynamics (Cabral et al., 2014; Deco, Cabral, et al., 2017).

Additionally, the WC model reveals frequency-specific FC (Figure 2a-b, i) and spatiotemporal dynamical patterns (Figure 3a-b, i) for a broader range of coupling strength and specific range of mean delay values when compared to the SL model (Figure 2-3, ii). This may be a consequence of the homeostatic mechanism in the WC model, which compensates for changes in coupling by regulating the local excitatory/inhibitory (E/I) balance. More specifically, within reasonable bounds, local homeostatic plasticity compensates for the higher levels of incoming excitatory input resulting from increased global coupling, thus maintaining the local dynamics at a desired level. Furthermore, homeostatic plasticity attenuates the effect of different node degrees in the connectivity graph, allowing for more uniform dynamics across the connectome.

The importance of delays in the WC model — with homeostatic plasticity — is even more apparent when analysing the fit to empirical FCD over global couplings and target firing rates (SM, Section I, Figure S5). Taking BOLD signals as an example: a mean delay of 4ms affords a broad region in parameter space where FCD is accurately represented, which is not seen for other mean delays. Accordingly, we clearly see the differential role of global parameters in tuning the underlying simulated network dynamics when a local parameter (i.e., firing rate) is considered, highlighting the importance of multiscale interactions. On the other hand, under the SL model, the coupling and the conduction delays gives rise to frequency-specific connectivity patterns: The further the frequency of interest is from the intrinsic frequency of the model, the greater the range of possible optimum delays (Figure 2b, ii). Interestingly, in the SL model, the best agreement (white stars) follows a frequency-suppression rule: the lower the frequency band in which the signal is filtered, the longer the range of optimal delays.

This suggests that in the presence of delays — and when nodes are sufficiently connected — the frequency of global connectivity decreases due to an increase in overall synchrony. Besides, for the WC model, FCD deteriorates for higher levels of coupling where the network engages in supercritical dynamics, abolishing the relationship between frequency suppression and synchrony at lower frequencies.

89 HCP participants) and simulated Hilbert envelope FCD histograms for each pair of parameters, for theta [4-8 Hz] (*left*), alpha [8-13Hz] (*middle*), beta [13-30Hz] (*right*) for WC and SL model. **i)** *Top –* For the WC model, the optimal points for MEG are C=0.34, MD=3ms for theta; C=0.42, MD=3ms for alpha; C=0.78, MD=4ms for beta with ks-distance values of ks_theta=0.084, ks_alpha=0.194, ks_beta=0.395. *Bottom -* Empirical and simulated frequency-specific MEG FCD distribution for 78 AAL cortical brain areas in the optimal point. **ii)** *Top -* For the SL model, the optimal points for MEG are C=549, MD=4ms for theta; C=711.32, MD=1ms for alpha; C=194.8, MD=2ms for beta with ks distance values of ks_theta=0.168, ks_alpha=0.104, ks_beta=0.149. Stars indicate the optimal point where both FC correlation and the FCD ks distance are maximised and minimised, respectively. *Bottom -* Empirical and simulated frequency-specific MEG FCD distribution for 78 AAL cortical brain areas in the optimal point. White stars indicate the model working points, chosen through simultaneous optimization for the reproduction of empirical FC and FCD, as described in the Methods section.

To assess the impact of integrating delays, quantitative results are also shown for the null-delay scenario (Tables 2 and 3). Interestingly, the role of delays is even more evident when optimising for each of the features individually (SM, Section II, Table S1-S2). Table 2 and Table 3 show the correlation and distance values for FC and FCD optimised simultaneously, respectively; in both cases, including and excluding delays.

**Table 2.** Performance values, optimised for both FC and FCD features, accounting for delays. Note that for Pearson correlation a greater value corresponds to a better fit whereas for KS distance it’s the opposite.

| Best fit (Pearson correlation) with delays, optimised for FC and FCD |  |  |  |  | Best fit (KS distance) with delays, optimised for FCD and FCD |  |  |  |
| --- | --- | --- | --- | --- | --- | --- | --- | --- |
|  | fMRI | MEG<sub>Theta</sub> | MEG<sub>Alpha</sub> | MEG<sub>Beta</sub> | fMRI | MEG<sub>Theta</sub> | MEG<sub>Alpha</sub> | MEG<sub>Beta</sub> |
| WC | 0.479 | 0.460 | 0.465 | 0.514 | 0.107 | 0.084 | 0.194 | 0.395 |
| SL | 0.469 | 0.471 | 0.494 | 0.588 | 0.489 | 0.168 | 0.104 | 0.149 |

**Table 3.** Performance values, optimised for both FC and FCD features, disregarding delays. Note that for Pearson correlation a greater value corresponds to a better fit whereas for KS distance it’s the opposite.

| Best fit (Pearson correlation) without delays, optimised for FC and FCD |  |  |  |  | Best fit (KS distance) without delays, optimised for FCD and FCD |  |  |  |
| --- | --- | --- | --- | --- | --- | --- | --- | --- |
|  | fMRI | MEG<sub>Theta</sub> | MEG<sub>Alpha</sub> | MEG<sub>Beta</sub> | fMRI | MEG<sub>Theta</sub> | MEG<sub>Alpha</sub> | MEG<sub>Beta</sub> |
| WC | 0.419 | 0.331 | 0.286 | 0.144 | 0.268 | 0.721 | 0.712 | 0.458 |
| SL | 0.458 | 0.226 | 0.217 | 0.443 | 0.584 | 0.074 | 0.147 | 0.149 |

There are a few features that can be identified from the ensuing distributions of FCD values. First, empirical BOLD distributions have a longer tail than the distributions for all MEG frequencies. This is likely due to the 80% overlap between sliding windows used for BOLD signals, which leads to higher correlation values close to the diagonal of the FCD matrix. Second, MEG distributions shift towards higher correlation values for higher frequency bands. Since we used the same window size (2s) for all frequencies, higher frequency oscillations lead to more cycles within a sliding window and, subsequently, to more fluctuations in amplitude. Correlations between signals are, therefore, less impacted by noise.

To summarise the ability of both models to reproduce static and dynamical fMRI/MEG patterns, we performed a cross-modality analysis to search for a region of conjunction (Figure 4). Regarding FC, as shown in the previous results, for the WC model there is a narrow vertical region where the delay must be very specific to ensure a good fit across modalities. Therefore, we find no consistent region where FC can be accurately represented for all modalities. However, this analysis confirms the importance of accounting for delays in neural mass modelling.

**Figure 4.**
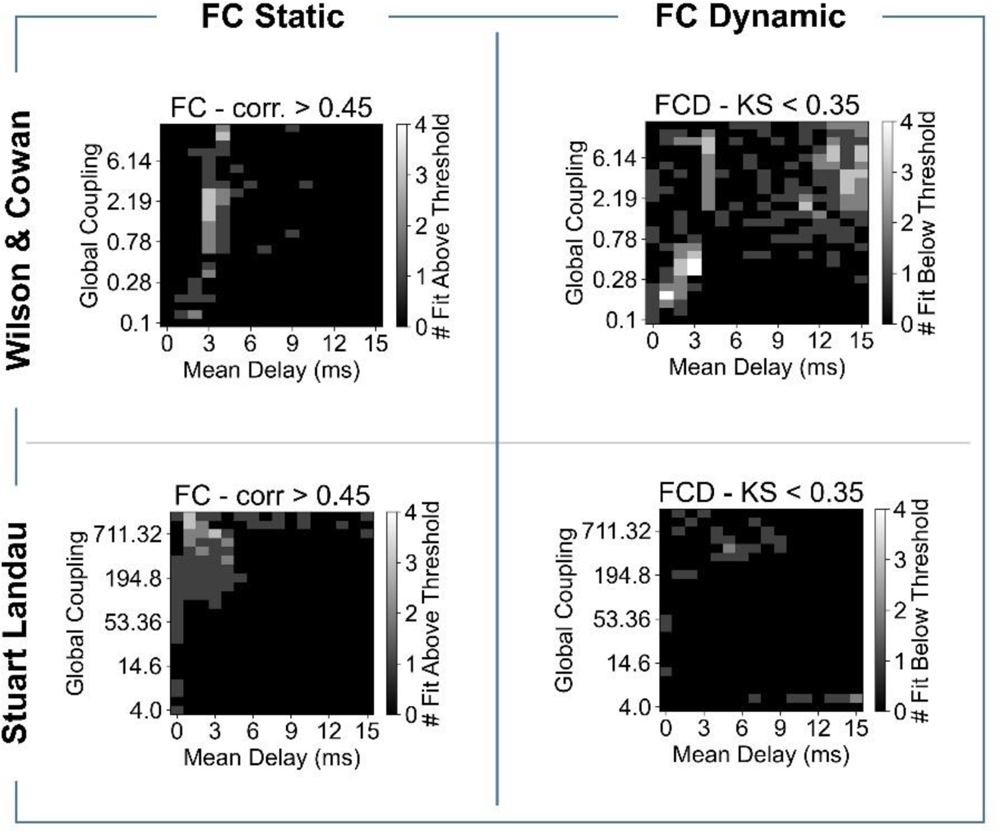
Model performance in explaining static and dynamical connectivity features across modalities. Agreement plots indicating the number of modalities (BOLD, MEG theta, MEG alpha, MEG beta) with correlation between empirical and simulated FC above 0.45 and KS distance below 0.35 for Wilson and Cowan and Stuart Landau model.

On the other hand, the SL model is generally better at simultaneously reproducing empirical FC across modalities. One plausible reason for this is that, for each modality, there is a broad region where FC can be adequately reproduced so it is easier to find regions of overlap. Although the more we move towards slow signals the more this region expands horizontally, there is an optimal range of parameters - that is between 1-4ms and for high couplings, where the model evinces an alignment in performances across modalities.

Regarding the FCD, the cross-modality performance of the SL gets considerably worse. There are no points in the parameter space where the model can perform well for more than two modalities simultaneously. Therefore, this result suggests that SL dynamics are more sensitive to changes in global parameters such as couplings and delays. For the WC model, this dialectic is not so evident. In fact, there is a region of fast delays and weak coupling where simulated FCD accurately represents empirical results across modalities.

Furthermore, it appears there is a stronger relationship between delays and couplings in shaping WC FCD, where longer delays require stronger couplings. The most likely reason the WC model performs better in this regard is the fact that local dynamics are actively regulated towards a common target across the brain, through E-I homeostasis. This reflects a multiscale interaction between local and global dynamics that renders WC dynamics less dependent on the global parameters.

### 3.2 Rich but constrained functionality emerges from the invariant anatomical architecture

The repertoire of functional networks lies upon the hidden structural architecture of connections that facilitates hierarchical functional integration (Park and Friston 2013). Here, we explore the performance of two large-scale generative models, with the goal of understanding the underlying processes that give rise to coherent large-scale functional networks. Nonetheless, both modelling approaches have the human DTI-based structural connectome as the only empirically derived element. Therefore, such models can also be understood as a nonlinear system which, taking the connectome as the input, can be used to evaluate the possible causal mechanisms for a phenomenon of interest to emerge (i.e., functional connectivity patterns). By investigating the non-trivial structure-function relationship, we highlight two main findings: simulated FC correlates with empirical FC better than the structural connectivity (SC) alone (Figure 5). However, this correlation is still lower than the correlation between simulated FC and SC (Figure 6).

**Figure 5.**
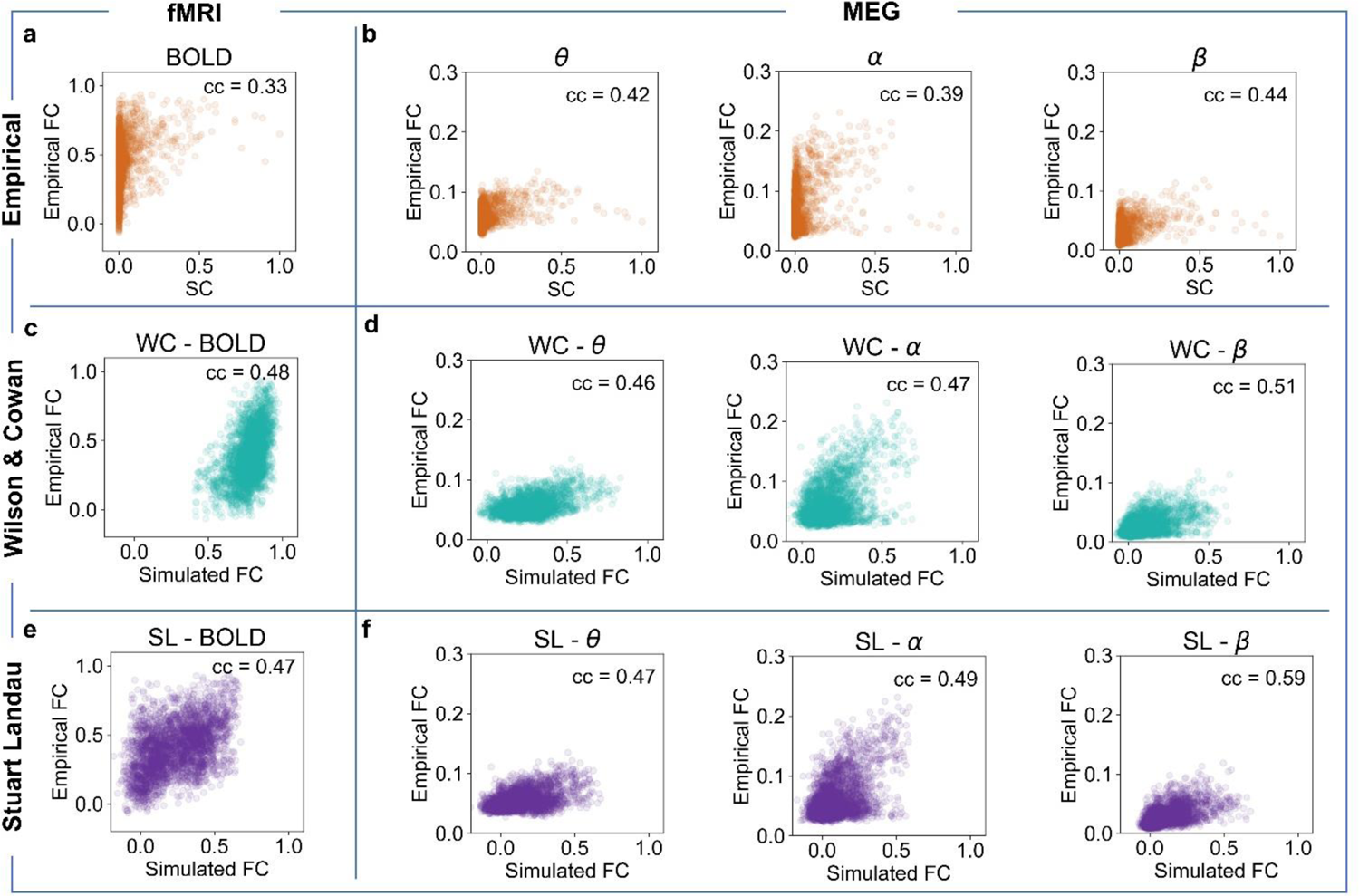
Beyond the anatomical framework. **a.** Scatter plot of Empirical Functional Connectivity (FC) versus empirical Structural Connectivity (SC) with a correlation value of cc=0.33. **b.** Scatter plot of MEG FC versus SC for theta (left), alpha (middle), beta (right) with a correlation value of cc_Theta_=0.42, cc_Alpha_=0.39, cc_Beta_=0.44. **c.** Scatter plot of BOLD fMRI FC versus simulated WC FC with a correlation value of cc=0.48. **d.** Scatter plot of empirical versus simulated WC MEG FC SC for theta (left), alpha (middle), beta (right) with a correlation value of cc_Theta_=0.46, cc_Alpha_=0.52, cc_Beta_=0.42. **e.** Scatter plot of Empirical BOLD fMRI FC versus Simulated SL FC with a correlation value of cc=0.47. **f.** Scatter plot of empirical versus simulated SL MEG FC for theta (left), alpha (middle), beta (right) with a correlation value of cc_Theta_=0.45, cc_Alpha_=0.49, cc_Beta_=0.59.

**Figure 6.**
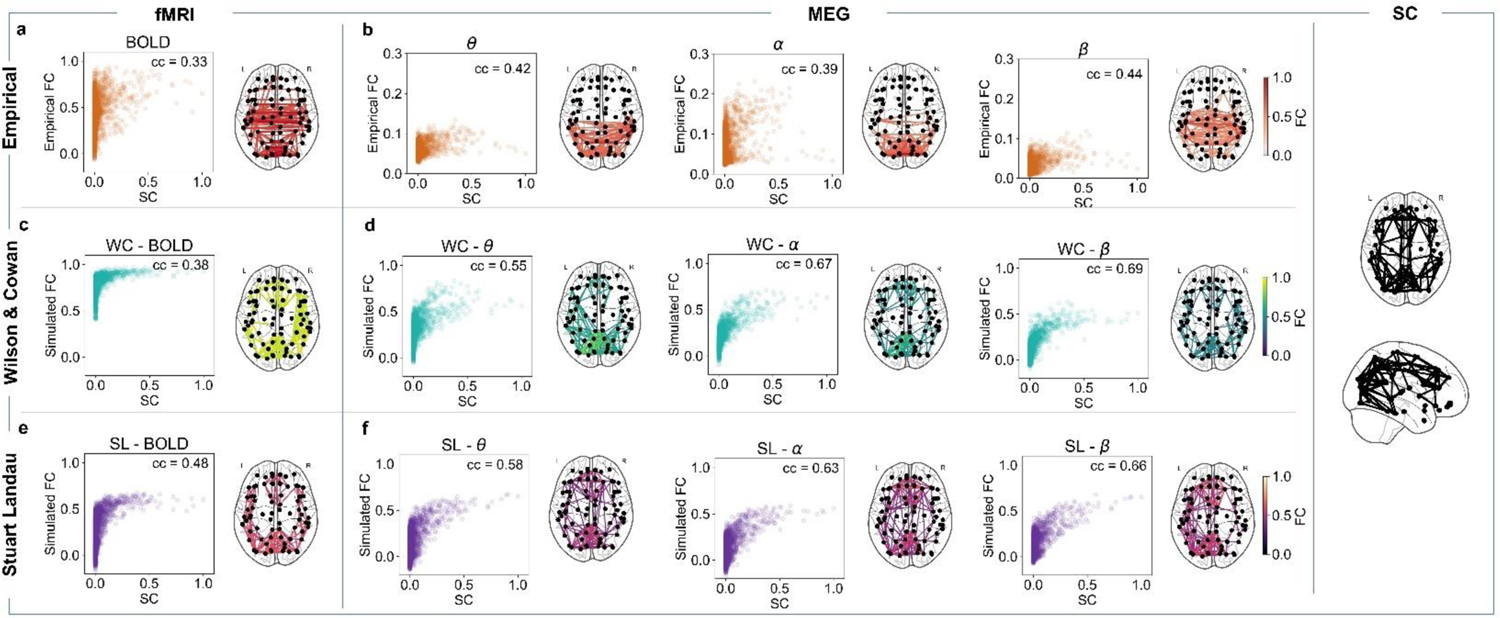
The role of the connectome. **a.** Scatter plot of empirical BOLD fMRI Functional Connectivity (FC) versus Structural connectivity (SC) with a correlation value of cc=0.33. **b.** *Left* - Scatter plot of MEG theta FC versus SC with a correlation value of cc=0.42. Note that the correlation plots here are the same as in Fig. 5 (a, b) and are replicated here for the reader’s convenience. *Middle -* Scatter plot of MEG alpha FC versus SC with a correlation value of cc=0.39. *Right -* Scatter plot of MEG beta FC versus SC with a correlation value of cc=044. Brain plots showing the 5% strongest connections of empirical FC. **c.** Scatter plot of simulated WC BOLD fMRI FC versus SC, with a correlation value of cc=0.38. **d.** *Left -* Scatter plot of simulated WC MEG theta FC versus SC, with a correlation value of cc=0.55. *Middle -* Scatter plot of simulated WC MEG alpha FC versus SC, with a correlation value of cc=0.55. *Right -* Scatter plot of simulated WC MEG beta FC versus SC, with a correlation value of cc=0.69. **e.** Scatter plot of simulated SL BOLD fMRI FC versus SC, with a correlation value of cc=0.48. **f.** *Left -* Scatter plot of simulated SL MEG theta FC versus SC, with a correlation value of cc=0.58. *Middle -* Scatter plot of simulated SL MEG alpha FC versus SC, with a correlation value of cc=0.63. *Right -* Scatter plot of simulated SL MEG beta FC versus SC, with a correlation value of cc=0.66. Brain plots showing the 5% strongest connections of simulated FC.

More specifically, figure 5 shows that, for both models, simulated FC is generally better than SC patterns alone at explaining empirical FC. This can be clearly seen in both BOLD fMRI and in the MEG cases, where the agreement between SC and simulated FC is significantly higher than between SC and empirical envelope FC, demonstrating the capability of generative models to model dynamics that the structure alone is unable to reproduce. Under this perspective, it is worth noting that the empirically derived input to our models is not limited to the structural connectome, but also the tract-length between each pair of areas. Therefore, while not impacting directly on the structure of functional interactions, the information from tract lengths is relevant, when including mean delays.

Concomitantly, large-scale models are useful to validate the predictive validity of structural information in relation to predicting the functional connectivity, proving that the realistic between-area strength of the SC is necessary for functional networks and their related topological features to emerge. As a proof of concept, without the right structure there is no emergence of function (SM, Section III, Figure S13).

Notwithstanding the benefit of the above-mentioned biophysical models, the correspondence between structure and function is still not optimal (cc_Max_ < 0.6). One of the possible reasons may be that the alignment between structural and empirical functional data is weak in both fMRI and MEG scenarios (Figure 6a-b). Additionally, as shown in Figure 6c/f, the functional topology patterns seem being largely constrained by structure, regardless of the model implemented — the relationship between FC and SC remains the same for both models.

### 3.3 Metastable oscillatory modes reflect coupling among brain regions

A key feature of the metastable regime is a dynamic modulation of the frequencies in the connectome expressed by the itinerant succession of dynamical modes. To address the temporal evolution of simulated and empirical signals we investigated how inherently unstable neural communities can engage in stable transient and partially synchronised modes, named metastable oscillatory modes (MOMs) (Cabral et al., 2022).

To detect MOMs and characterise them in space and time, we band-pass filter the simulated and empirical signals around the peak frequency and obtain the corresponding amplitude envelopes using the Hilbert transform. Here, we consider that a brain region evinces a MOM if the amplitude increases by 2 standard deviations above the global mean amplitude in that frequency range (coloured shades in Figure 7) (see *Metastable Oscillatory Modes*, Methods). We illustrate MOMs for 800s of BOLD fMRI signals, and 10s of source reconstructed MEG signals. To demonstrate their concurrent appearance, we display simulated MOMs in one single point of the parameter space for each model. To choose this point, for each model, we pick the parameters for which the model best describes functional connectivity dynamics across modalities (SM, Section II, Figure S9). Nonetheless, models can generate MOMs with similar properties in other points of the parameter space when delays are considered (SM, Section II, Figure S12). As shown in figure 7, all areas engaging in each coalition exhibit the simultaneous emergence of an oscillation, resonating at the same collective frequency.

**Figure 7.**
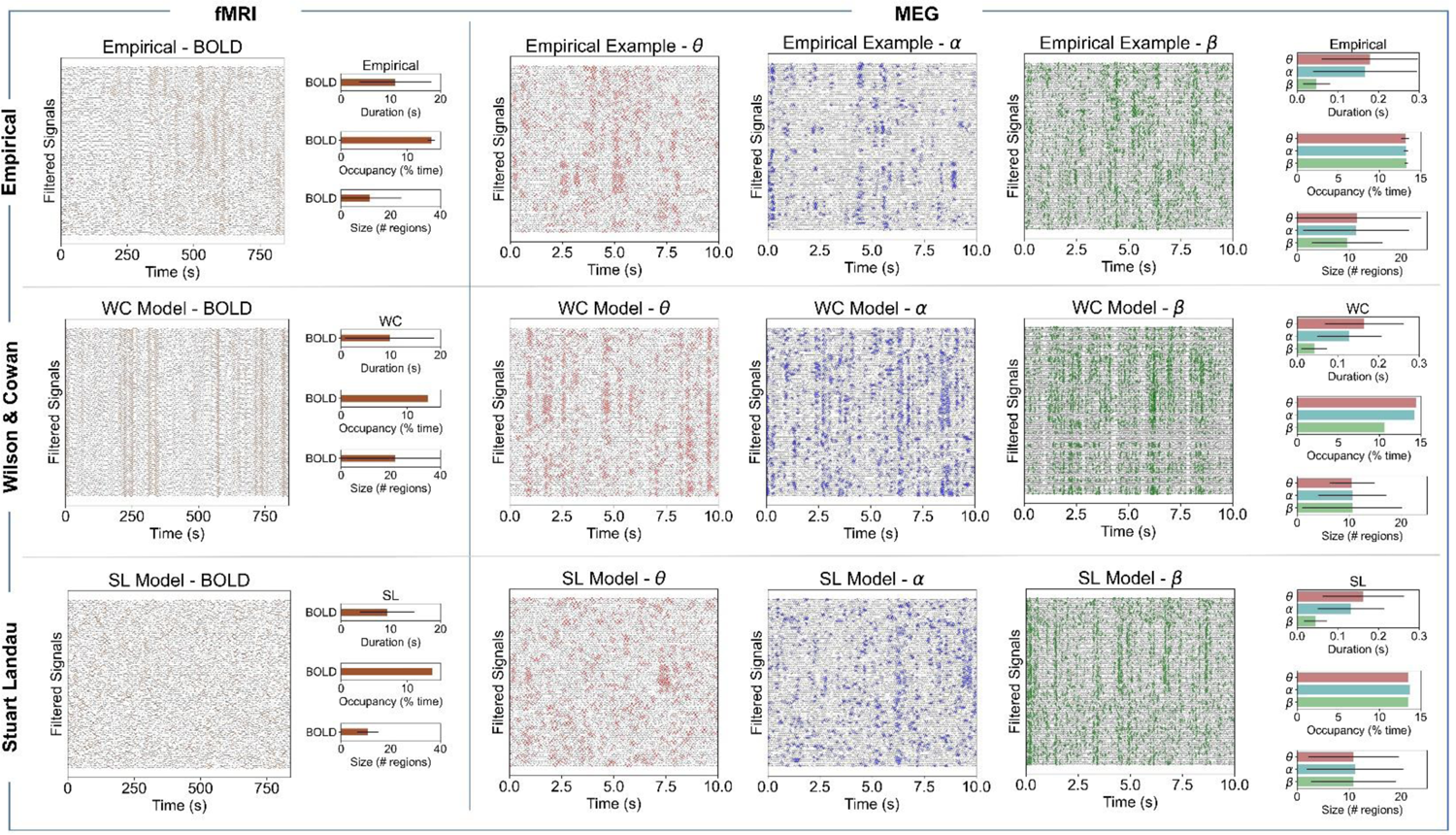
Emergence of transient-like dynamics in empirical and simulated signals. Examples of empirical and simulated signals in 78 anatomically-defined regions plotted over 800s minutes for BOLD fMRI and 10 seconds for MEG. The coloured parts show the points in time when the signal power exceeds a certain threshold. For each modality, the threshold is defined as the 2-standard deviation of the amplitude of the signal itself. The transient itinerance is characterised in terms of duration, occupancy, and size (see Methods).

This analysis reveals distinct spatially-organised subsystems, which are thought to contribute to the formation of functional connectivity maps. More precisely, we find that both simulated and empirical MOMs are structured in space and emerge simultaneously across several brain areas, but also in time, here lasting about 10s for BOLD, between 100 and 200ms for MEG alpha and MEG theta, and less than 100ms for MEG beta.

## 4 Discussion

MEG and BOLD signals are believed to reflect two different aspects of neural activity, occurring at timescales that are orders of magnitude apart. While BOLD signals are thought to represent changes in haemodynamics (Hillman, 2014) — likely triggered by synaptic transmission; namely, the most energy intensive process in the human brain (Harris et al., 2012) — MEG signals reflect changes in magnetic fields created by dipole currents that flow along neuronal processes (Lopes da Silva, 2013). These dipole currents depend on dendritic synaptic input (Lopes da Silva, 2013) and are, therefore, related to the same processes involved in the generation of BOLD signals. Nonetheless, even though both modalities share the same neural substrate, large-scale models, to date, are usually tailored to represent only one modality at a time. A common example is the practice of tuning the intrinsic frequency of oscillation of local populations to the frequencies of interest in the respective modalities (e.g. ∼10 Hz for MEG, <0.01 Hz for BOLD) (Abeysuriya et al., 2018; Deco, Cabral, et al., 2017; Deco, Kringelbach, et al., 2017). In this work, we argue that models should be able to simultaneously generate multiresolution modalities with the same underlying generative (neuronal) mechanisms, without tuning parameters *a priori* to selectively reproduce features of interest. Combining multiresolution, multimodal data with large-scale modelling allows one, potentially to disentangle the generative mechanisms behind brain function and its dynamical underpinnings.

The first step we take towards cross-modality is to impose an intrinsic oscillation frequency of ∼40Hz, in the gamma range. In the human cortex, gamma rhythms are thought to be generated by a myriad of mechanisms (Buzsáki & Wang, 2012), such as reciprocal interactions between pyramidal neurons and fast-spiking interneurons (Buzsáki, 2006). Indeed, BOLD signal fluctuations have been hypothesised to arise from changes in synchrony between oscillators in the gamma band (Deco et al., 2009). Furthermore, in the case of MEG, recent results suggest that functional networks in lower-frequency bands (i.e. theta, alpha and beta) can be generated through delayed interactions between gamma oscillators (Cabral et al., 2022). Therefore, multiresolution recordings might reflect different aspects of gamma activity and models with local gamma oscillations might reproduce the empirical properties of both BOLD and MEG FC.

Models of brain networks should not only reflect the statistical dependencies among brain areas (i.e. FC), but also the dynamics underlying the spontaneous and transient appearance of functional networks (Baker et al., 2014; Cabral et al., 2017; Quinn et al., 2019; Vidaurre et al., 2016; Vohryzek et al., 2020). Therefore, we focus on the performance of models in the representation of both FC and its dynamics (FCD) across modalities. As demonstrated by recent research (Deco, Kringelbach et al. 2021 this approach is relevant not only because it offers more constraints for model validation, but also because FCD has been tied to the metastable dynamics characteristic of healthy brain function (Deco, Kringelbach et al. 2017).

### 4.1 Role of delayed interactions

Conduction delays have been shown to define a rich dynamic framework for the emergence of resting brain oscillations (Abeysuriya et al., 2018; Cabral et al., 2014; Petkoski & Jirsa, 2019). Building on previous research (Cabral et al., 2022), by including delays, we see a significant improvement of model performance when explaining empirical-wise MEG static and dynamic patterns.

However, in the fMRI context, while the WC generates patterns similar to empirical patterns for both cases (with and without delays), the SL model is sensitive to changes in delays. We think that this decreased dependence of low frequency oscillatory patterns on the mean delay (for the explored range) is related to the ratio between the period of oscillations and the delay itself. For example, while for beta rhythms (∼25 Hz, 40ms period) a change of 10ms in the mean delay represents the 25% of an oscillation cycle, for the slower BOLD rhythms (∼0.01 Hz, 100s period), the same change only accounts for 0.01% of a cycle. Therefore, we hypothesise that, for higher frequency bands, changes in conduction velocity have a greater impact in the ability of regions to synchronise at those frequencies due to greater changes in phase-relationships.

On this note, our results highlight the importance of delays for the generation of slow signals, such as BOLD fluctuations, when implementing neural mass models with “synaptic-like” communication between nodes (i.e., WC models). Accordingly, previous research implementing similar models informed by a macaque connectome (Deco et al., 2009), showed that BOLD signal fluctuations could be generated by transient synchronisation of coupled Wilson-Cowan nodes resonating at 40Hz. Importantly, this model was also sensitive to changes in conduction velocity, showing an optimal range of conduction speeds, even though local E-I balance was not modelled. Moreover, previous modelling results suggest that deficits in the regulation of axonal myelination could have a significant impact on the ability of coupled oscillators to synchronise at high frequencies, as our results suggest as well (Pajevic et al., 2014). Finally, the profound importance of modelling conduction delays was established using Bayesian model comparison (comparing models with and without delays) at the inception of dynamic causal modelling for fast, event-related responses as measured with EEG (David et al., 2006).

The effect of conduction delays was not seen for the SL model in the context of fMRI, likely due to the fact that the coupling is diffusive, rather than synaptic-like, thus reflecting the phase relationships between oscillations at specific frequency bands. Therefore, as mentioned above, the same interaction delay can have a different impact on oscillations with frequencies that are orders of magnitude apart. As a whole, these results speak to the use of delayed interactions, especially when the purpose is to find a common explanation for static and dynamic features of different neuroimaging modalities.

The importance of modelling delays leads to questions regarding their role in the brain. Our results suggest that conduction delays underscore the emergence of relevant dynamics, especially when looking at frequency-specific oscillatory bands (high-frequency in the SL, across frequencies for the WC). Therefore, it would follow that axonal conduction velocities should be precisely structured in the human brain. However, empirically, there is a high level of heterogeneity in the distribution of axonal diameters and levels of myelination (Boshkovski et al., 2021; Liewald et al., 2014; Lutti et al., 2014; Pajevic et al., 2014), both of which determine conduction speeds (Powanwe & Longtin, 2019; Saab & Nave, 2017), with complex interactions between both (Waxman, 1980). Furthermore, research suggests a dynamical regulation of myelination, at least in sensory systems (Saab & Nave, 2017). We suggest that such heterogeneities (or activity -dependent myelination) are an important aspect of computational architectures and message passing in the brain. Accounting for heterogeneous conduction velocities could help large-scale models — such the ones implemented here — to better explain empirical patterns of MEG connectivity, which are less clearly constrained by structural connectivity.

### 4.2 Role of E/I balance in the WC model

The importance of excitatory-inhibitory balance for cortical function is well known in the literature (Dehghani et al., 2016; Froemke et al., 2007; Sprekeler, 2017; Tao & Poo, 2005; Xue et al., 2014), along with the existence of synaptic plasticity in response to perturbations and developmental changes (Ma et al., 2019; Turrigiano, 2011; Turrigiano et al., 1998; Vogels et al., 2011).

Furthermore, the implementation of an extended version of the WC approach, builds on previous and established work demonstrating the significance of accounting for E-I balance in neural-mass models with excitatory and inhibitory populations (Abeysuriya et al., 2018; Deco et al., 2019; Hellyer et al., 2016).

One of the first effects - observed with the addition of the inhibitory synaptic plasticity (Abeysuriya et al., 2018; Vogels et al., 2011) is the tolerance of the model at strong coupling levels (SM, Section I, Figure S4). This tolerance allows one to explore a wider region of parameter space. Furthermore, since our cortical connectome has a wide range of node degrees (sum of incoming connections to a node) — which vary by at least one order of magnitude — nodes receive varying levels of excitatory input. Therefore, including a compensatory mechanism to adjust local dynamics protects against these discrepancies and generates more uniform dynamics across the brain.

Accordingly, our results show that the ability of the WC model with plasticity to accurately reproduce FCD features is less sensitive to parameter variations than the SL, particularly with respect to BOLD signals. This suggests that E-I homeostasis has a strong impact, especially on slow dynamics. For example, while adding plasticity does not impact greatly on the ability of the WC model to represent MEG FC and FCD, the same is not seen for BOLD signals, with models unable to represent BOLD FC when homeostasis is not modelled (FC cc = 0.2, FCD KS = 0.13) (SM, Section I, Figure S4). In addition, while the WC model without plasticity can reproduce MEG accurately, it does so for combinations of parameters with a broader range of mean delays (BOLD = 8ms, MEG_Theta_ = 5ms, MEG_Alpha_ = 2ms, MEG_Beta_ = 0 ms), compared to the WC model with plasticity (3-4 ms). Therefore, we suggest that the addition of E-I homeostasis is important for cross-modality validity of generative models, particularly regarding the reproduction of ultra-slow dynamics.

### 4.3 Structure-function relationship

One of the most important issues when modelling large-scale brain activity – with a structural connectome as a substrate — is to ensure that the models with dynamics outperform models based on the structural connectome alone. By contrasting the correlation between empirical FC and SC vs empirical FC and simulated FC, our results show that for both models and modalities, the optimised models with dynamics always provide an improvement in the prediction of empirical FC. Furthermore, this difference is even more evident when optimising models to represent FC only (SM, Section II, Table S1-S2). Therefore, both models used to model local dynamics, Stuart-Landau and Wilson-Cowan, account for emergent properties of human FC that are not captured by structural connectivity alone. Besides stressing the role of non-linear dynamics and interactions in brain networks, this further establishes the validity of integrating delayed interactions in models for the prediction of even static FC (Cabral et al., 2011; Deco et al., 2009). In fact, although there is an established exponential relationship between connection strength and distance (Ercsey-Ravasz et al., 2013), there are exceptions to this rule, shown to be relevant for large-scale functional networks (Gustavo Deco et al., 2021). Therefore, we argue that the conduction delays between areas enrich models beyond the underlying structural framework.

Furthermore, although the WC model with plasticity performed better when reproducing dynamical spatiotemporal features (FCD) especially for BOLD, its added complexity did not guarantee a better resemblance of functional patterns. This suggests that models could benefit from the inclusion of more detailed empirically-derived information about sources of heterogeneity such as local microcircuitry (Wang, 2020) or myelination (Boshkovski et al., 2021) as discussed in detail below.

Importantly, a common finding — across modalities and modelling approaches — is the higher correlation between simulated FC and SC, compared to empirical FC and SC. This leads to two possible and not exclusive interpretations: First, both modelling approaches are overly constrained by the structural connectome that lacks information about the strength of effective connectivity and the direction of connectivity (e.g., forwards versus backwards). In addition, research suggests that there are gradients in microcircuitry organisation, such as asymmetries in laminar specific forward and backward connections and recurrent excitation or the distribution of inhibitory interneurons (Wang, 2020), that reflect the hierarchical organisation of the human cortex (Felleman & Van Essen, 1991). Not only is this hierarchical organisation functionally relevant for processes such as perception (van Vugt et al., 2018) and memory (Froudist-Walsh et al., 2021), but recent modelling results show that accounting for these asymmetries improves the reproduction of FC and FCD, while allowing for the emergence of important (i.e., non-dissipative) dynamics, such as vortices, turbulence and ignition dynamics (G. Deco et al., 2021). In addition, the spatial distribution of such asymmetries and variations in synaptic time constants might also explain why we observe particular frequency bands more strongly in certain anatomical regions, as is the case of beta in the parietal cortex and alpha in the occipital lobe. Indeed, myelination imaging indicates that these regions include areas with the highest myelin content (Matthew F. Glasser et al., 2016; Rowley et al., 2015), which could relate to higher conduction speeds (Saab & Nave, 2017), favourable to the emergence of relevant functional networks at these higher frequencies.

Second, structural connections may be underestimated using tractography. One example is the limited ability of DTI to estimate interhemispheric white matter tracts, leading to a difficulty in reproducing the strong homotopic interhemispheric functional correlations present in fMRI (Deco, Ponce-Alvarez, et al., 2013). Furthermore, recent results show that communication between cortical areas at different frequency bands has varying degrees of dependence on the underlying anatomy (Vezoli et al., 2021), suggesting that empirical FC reflects processes that go beyond structure; i.e., functional connectivity reflects ‘dynamics on structure’.

### 4.4 Metastable Oscillatory Modes

Analysis of recurrent metastable oscillatory modes may elucidate the mechanisms behind the functional integration - segregation relationship (Friston, 1997; Friston, 2000). As phenomenologically shown in recent work (Cabral et al., 2022), and further validated in the current study, when the coupling is sufficiently strong, the emergent dynamics will start to resemble the complex and intermittent dynamics observed in neuronal timeseries. As we further increase the extrinsic coupling of our models, the system locks into a regime of complete entrainment losing the frequency-specific intermittency.

In both models, we selected a single point in parameter space to illustrate how the concomitant emergence of frequency-specific coherent patterns manifests, from slow to fast oscillations. This speaks to a unified model of the brain - in which each parameter combination can reflect a particular neural state with its own prominent frequency, specific connectivity, and network topology. Remarkably, in this study, MOMs were detected in both fMRI and MEG signals. Although similar methods have been successfully applied in the context of fast oscillations (i.e. MEG (Sorrentino et al., 2021)) the appearance of metastable fluctuation in the fMRI context is relatively unexplored.

Our results suggest that self-limiting transient oscillations are detectable also in signals with spontaneous sustained periodicity, such as fMRI timeseries. This is in line with the notion that synchronisation underscores fMRI correlations (Lu et al., 2007) and the potential of fMRI to map neural oscillations (Lewis et al., 2016), suggesting the possible coexistence of both transient events and sustained oscillations in the brain (van Ede et al., 2018). Furthermore, characterising MOMs according to properties such as size, occupancy and duration can help in validating the dynamic aspect of the models and investigating similarities with empirical data.

## 5 Limitations and Future Work

### 5.1 Averaging (over subjects)

We used the structural connectome - derived from the average of 32 DTI scans - in this work to define the connectivity matrix. These data were acquired as part of a study separate from the MEG and fMRI HCP data. Averaging over subjects in DTI studies is deemed a necessary step in order to reduce the effect of signal loss due to changes in local magnetic susceptibility, which can lead to the aberrant inferences about diffusion direction being estimated and false positives and false negatives (Damoiseaux & Greicius, 2009).

In effect, we used the average structural connectivity matrix derived from one group to reproduce functional data similar to another group. We suspect that this may have limited our ability to find better correlations between the real and synthetic FCs. This issue suggests a similar analysis, in the future, where an individual’s tractography image is used to predict that subject’s MEG and fMRI features. In order to leverage the improved SNR of group-average data while accommodating heterogeneity over subjects (Quinn et al., 2021; Wens et al., 2014), a hierarchical model could be entertained.

### 5.2 MEG source reconstruction

Beamformers are a popular method for source reconstruction within the field of MEG, and have been used in FC studies (e.g. (Baker et al., 2014; Brookes et al., 2011; Hipp et al., 2012; Liuzzi et al., 2017). Often, they are chosen because of their ability to suppress sources of interference from outside source space (Boto et al., 2021; Cheyne et al., 2007; Litvak et al., 2010).

Despite their simplicity and popularity, beamformers are limited in the sense that they are, fundamentally, a spatial filter and therefore lack a generative model. This can make comparisons between alternative source inversion results non-trivial. Moreover, beamformers are known to suppress brain areas which exhibit high areas of zero-phase-lag (instantaneous) connections, i.e. correlated sources (Van Veen et al., 1997). Recent work has shown that using a beamformer for studies into the default mode network (DMN) at rest can be pernicious (Sjøgård et al., 2019). This provides an argument for using a source inversion algorithm with a full generative (i.e., forward) model which can account for correlations between brain areas in the source space, e.g. CHAMPAGNE (Owen et al., 2012) or Multiple Sparse Priors (MSPs) (Friston et al., 2008). However, at the time of writing, MSPs has been primarily optimised for time-averaged data and cannot readily be applied to resting-state scans.

An alternative approach - that we could have adopted in this work - would have been to side-step the ill-posed inverse problem altogether and instead focus efforts on maximising the similarity between sensor level covariance matrices (or some other statistic) of the simulated and real MEG datasets. This would have removed the confound of source leakage during the model screening process, although we would have to have accounted for variations in head position and greater levels of sensor noise which the beamformer implicitly reduces.

### 5.3 Generation of hemodynamic and electrophysiological data

One of the main limitations of our modelling approaches is the fact that while we used a generative approach to go from neuronal activity (e.g. LFPs or population firing rates) to BOLD signals (Buxton et al., 1998; Friston et al., 2000) we do not follow the same approach for generating MEG signals. Instead, we assume that the signals generated by our models can be directly mapped to source-reconstructed MEG. However, the MEG/EEG inverse problem is insoluble, and all source inversion algorithms (beamformers, minimum norm etc.) impose some form of assumption. In the fMRI context, hemodynamic models reflect the physiological relationship between population activity and the blood oxygenation measured through BOLD signals and they have been extensively validated (Buxton et al., 1998; Friston et al., 2000; Handwerker et al., 2012). Therefore, our results would benefit from a similar generative model to compute the source dipole currents that are detected via MEG (Lopes da Silva, 2013). Nonetheless, since both our models can still reveal empirically relevant spatiotemporal patterns of MEG signals in a comparable manner, one might argue that this issue undermine our conclusions.

### 5.4 Subcortical Structures

In this work, network dynamics are modelled without accounting for the influence of subcortical nodes. The first reason is due to inadequate subcortical resolution offered by common atlases used in our modelling (i.e., AAL, Schaefer, Desikan-Killiany). The second is related to the difficulty in modelling the dynamics of some subcortical structures using the SL and WC models, which either consider nodes as oscillators or as networks of reciprocally coupled excitatory and inhibitory neurons, suitable for cortical dynamics. While this approach could still be valid for structures such as the hippocampus, similar in structure to the cortex (Kandel, 2021), it would fail to accurately represent the dynamics of areas such as the striatum, which is mostly composed of inhibitory neurons (Lanciego et al., 2012), or the cerebellum, which has a distinct microcircuitry (Voogd & Glickstein, 1998). The omission of subcortical structures could impact our results, for example by disregarding the influence of widespread thalamocortical projections in the establishment of alpha rhythms (Halgren et al., 2019; Roux et al., 2013) and in supporting interhemispheric connectivity (Teipel et al., 2009; Wang et al., 2019). Nonetheless, such approaches would require more complex models with multilevel structures (Meier et al., 2022). See (van Wijk et al., 2018), for a fuller discussion of this issue in neural mass modelling.

### 5.5 E-I Homeostasis

Regarding the implementation of E-I homeostasis, we modelled E-I balance through inhibitory plasticity (Abeysuriya et al., 2018; Deco et al., 2019; G. Deco et al., 2021; Vogels et al., 2011). While research shows the importance of inhibitory connections for the maintenance of balance (Luz & Shamir, 2012; Vogels et al., 2013; Vogels et al., 2011), there are other mechanisms in place such as scaling of recurrent excitation (Turrigiano et al., 1998) or regulation of intrinsic excitability of excitatory populations (Desai et al., 1999), which have not yet been explored in large-scale models. While the oscillatory dynamics of WC nodes are determined by the excitatory and inhibitory time constants (see *Neural mass model,* Methods), changes in local inhibition might further affect local dynamics, especially in highly connected nodes, which require stronger local inhibition. Therefore, including additional homeostasis mechanisms, that synergistically interact with each other, may reveal relevant patterns of local microcircuitry, possibly related to gradients in cortical organisation (Wang, 2020).

### 5.6 Model Optimisation

On a more methodological level, the use of a grid search for model optimization, despite being common in large-scale modelling research (Cabral et al., 2022; Deco, Cabral, et al., 2017; Hellyer et al., 2016), is an inefficient method to explore the parameter space. This can be solved by making use recent advances such as Bayesian optimization (Hadida et al., 2018) and corresponding variational procedures used in dynamic causal modelling (Frässle et al., 2017; Razi et al., 2017). In addition, different metrics of performance could have been used to compare empirical and simulated data, such as power-spectrum similarity (Verma et al., 2022), and distance measures such as KL-divergence, KS-distance or mean-squared error between matrices (Savva et al., 2019).

### 5.7 Relationship between BOLD and MEG signals

One of the main perspectives offered by exploring the model performance across modalities is the fact that our models can generate simultaneous MEG and BOLD signals. This is relevant, given the fact that the relationship between MEG and fMRI signals is not yet fully understood (Garcés et al., 2016; Hall et al., 2014). In addition, recent results suggest that this relationship is not homogeneous across the brain, and that it is driven by differences in local circuitry related to the cortical hierarchy (Shafiei et al., 2022). Therefore, multimodal models might help to elucidate the interactions between the processes behind the two signals, particularly with studies involving the perturbation of dynamics through the application of external currents. We propose future studies to focus on the mechanistic relationship between MEG and fMRI, and how MEG features such as the relative power at different frequency bands, cross-frequency interactions and synchronisation can reflect the properties of hemodynamic signals. Please see (Friston et al., 2019; Jafarian et al., 2020; Wei et al., 2020) for further discussion.

### 5.8 Model Augmentation with Heterogeneity

Given our conclusions on the constraints imposed by the connectome in both models we explored, a crucial future step in modelling research is the inclusion of empirically derived sources of heterogeneity in large-scale computational models. Recent endeavours have shown the use of including transcriptomically derived differences in the excitability of local populations in the representation of static (Demirtaş et al., 2019) and dynamic (G. Deco et al., 2021) features of large-scale brain activity. In addition, results suggest that the variations in structure-function coupling across the cortical hierarchy are shaped by heterogeneities in local E-I balance and myelination levels (Fotiadis et al., 2022), or in cortico-subcortical interactions in terms of neuroreceptors density maps (Beliveau et al., 2017), temporal time-scales (Baldassano et al., 2017), gene expression (Hawrylycz et al., 2012), myelin content (in terms of T1/T2-weighted MRI signal) (Glasser & Van Essen, 2011) and functional connectivity (Kong et al., 2021) - offering further explanations as to why empirical FC exhibits characteristics that cannot be explained solely by SC. Therefore, we believe that it is essential for further modelling studies to make use of multilevel datasets (Arnatkeviciūtė et al., 2019; Royer et al., 2022) to constrain models with directed connectivity that define cortical hierarchies (G. Deco et al., 2021).

## Author contributions

Conceptualization: FC, FPS, RT, JC, VL Methodology: FC, FS, RT, JC, JV, VL, MW Investigation: FC, FS Visualization: FC, FS Funding acquisition: VL, KF, GD, PV Project administration: FC, VL Supervision: VL, KF, GD, JC, PV Writing – original draft: FC, FS Writing – review & editing: FC, FS, VL, KF, JC, JV

## Data and materials availability

All simulations and analysis were performed in Python except for the Source Reconstruction algorithm which was performed in MATLAB2021b. The codes and materials used in this study are available at: https://gitlab.com/francpsantos/whole_brain_generative_models

## Acknowledgments

We would like to thank Gareth Barnes and his team at the University College London, and the Oxford Centre for Human Brain Activity for the relevant insights and fruitful discussions. F.C. and F.P.S. are funded by the EU-project euSNN European School of Network Neuroscience (MSCA-ITN-ETN H2020-860563). The Wellcome Centre for Human Neuroimaging is supported by core funding from Wellcome [203147/Z/16/Z] and this funds F.C., R.T., V.L. and K.F. J.C. is funded by the Portuguese Foundation for Science and Technology grants UIDB/50026/2020,UIDP/50026/2020 and CEECIND/03325/2017. J.V. is supported by the EU H2020 FET Proactive project Neurotwin grant agreement no. 101017716. M.W. is supported by NIHR Oxford Health Biomedical Research Centre, the Wellcome Trust (106183/Z/14/Z and 215573/Z/19/Z), and the New Therapeutics in Alzheimer’s Diseases (NTAD) study supported by the MRC and the Dementia Platform UK. P.V. is founded by Virtual Brain Cloud (H2020 ID 826421). G.D. is supported by the Spanish national research project (PID2019-105772GB-I00 MCIU AEI), funded by the Spanish Ministry of Science, Innovation and Universities (MCIU), State Research Agency (AEI); HBP SGA3 Human Brain Project Specific grant agreement 3 (945539), funded by the EU H2020 FET Flagship program; SGR Research Support Group (reference 2017 SGR 1545), funded by the Catalan Agency for Management of University and Research Grants (AGAUR); Neurotwin Digital twins for model-driven non-invasive electrical brain stimulation (grant agreement 101017716), funded by the EU H2020 FET Proactive program; euSNN (grant agreement 860563), funded by the EU H2020 MSCA-ITN Innovative Training Networks; The Emerging Human Brain Cluster (CECH) (001-P-001682) within the framework of the European Research Development Fund Operational Program of Catalonia 2014–2020; Brain-Connects: Brain Connectivity during Stroke Recovery and Rehabilitation (201725.33), funded by the Fundacio La Marato TV3; and Corticity, FLAG-ERA JTC 2017 (reference PCI2018-092891), funded by the MCIU, AEI. Human data were provided by the Human connectome Project, WU-Minn Consortium (Principal Investigators: David Van Essen and Kamil Ugurbil; 1U54MH091657) funded by the 16 NIH Institutes and Centers that support the NIH Blueprint for Neuroscience Research; and by the McDonnell Center for Systems Neuroscience at Washington University.

