## Supplementary Methods for "Multi-modal and multi-model interrogation of large-scale functional brain networks"

### SECTION I

#### Uncoupled Node Dynamics

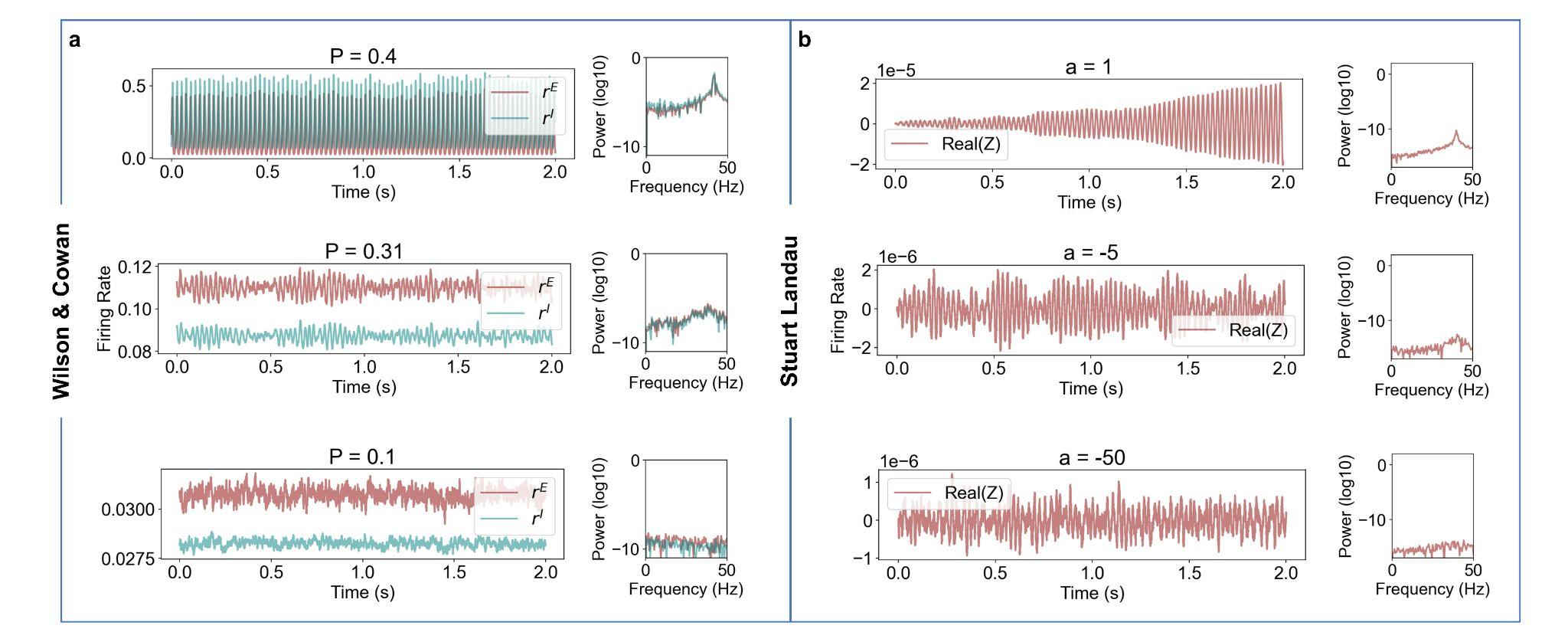

**Figure S1. Uncoupled node dynamics for the Wilson & Cowan and Stuart Landau models, at different values of the bifurcation parameter and fundamental frequency of 40Hz.** Each dynamical Wilson & Cowan (**a**) and Stuart Landau (**b**) oscillatory unit is perturbed with noise, where, depending on the value of the bifurcation parameter (*P* and *a* respectively), the model generates a pure oscillatory signal (top), oscillations with fluctuating amplitude (middle), noisy signal (bottom).

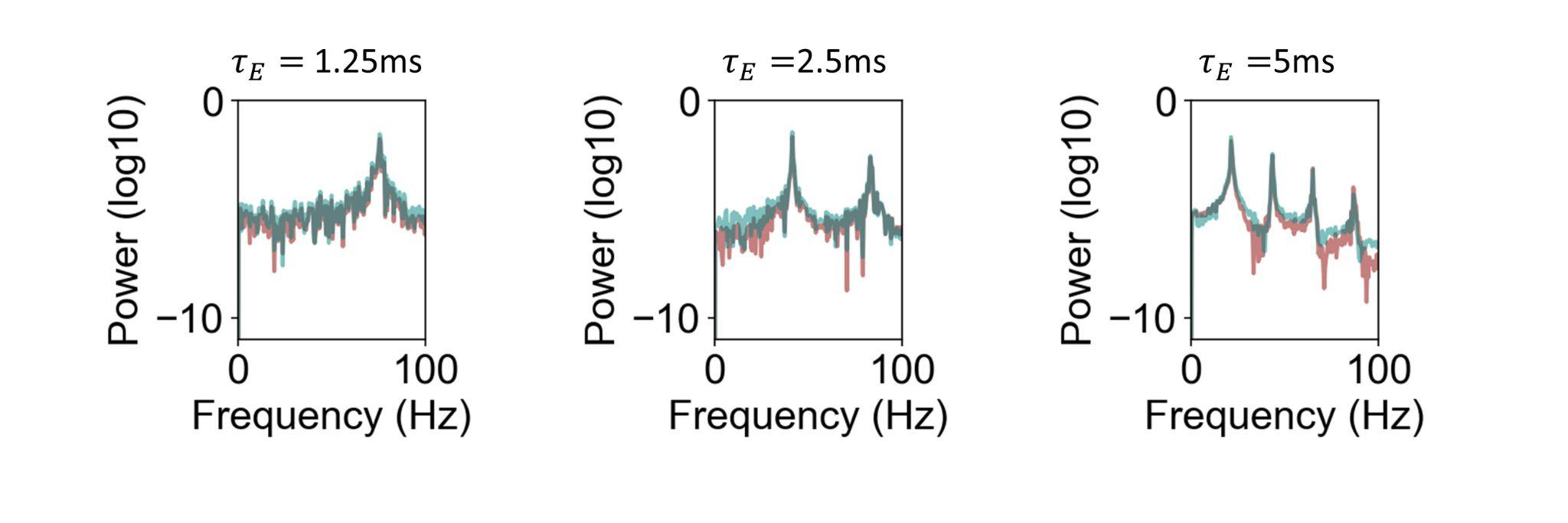
**Figure S2. Frequency profile for different values of** $\tau_{\boldsymbol{E}}$ **in the WC model.** Effect of the population time constants on the frequency of oscillation, where $\boldsymbol{\tau}_{\boldsymbol{I}}$=$2* \boldsymbol{\tau}_{\boldsymbol{E}}$. In this work we choose $\tau_{\boldsymbol{E}}$ and $\tau_{\boldsymbol{I}}$so that the characteristic frequency of isolated neural masses is within the gamma range (~40 Hz).

#### Homeostatic Plasticity

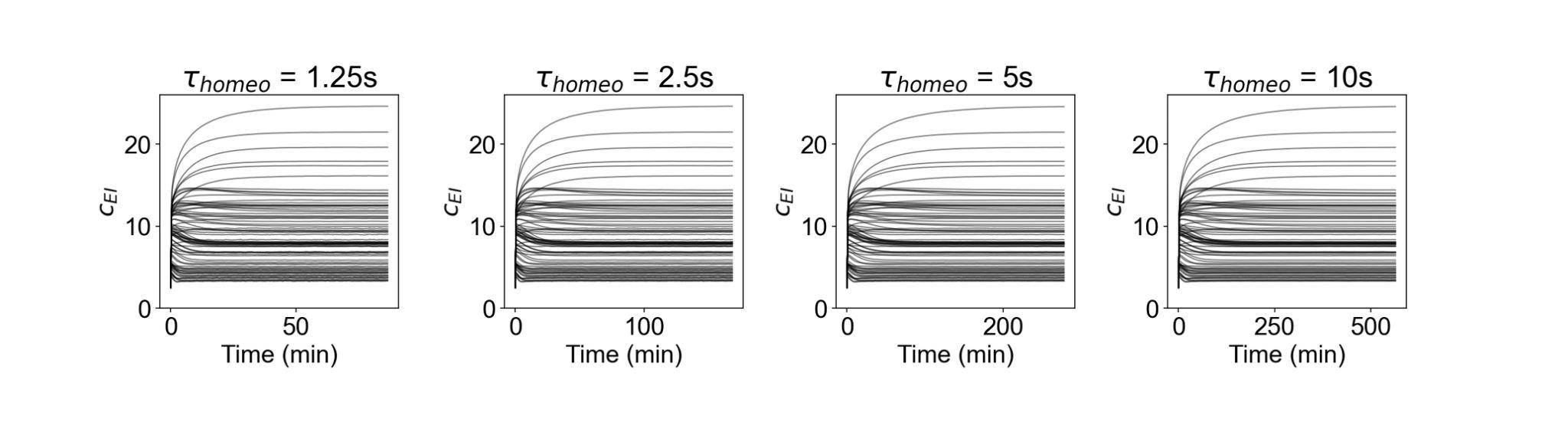

**Figure S3. Influence of the homeostatic plasticity on** $\boldsymbol{c}_{\boldsymbol{EI}}$**.** When homeostatic plasticity ($\tau_{homeo}$) is sufficiently slow to be decoupled from fast dynamics of intrinsic oscillations, $\boldsymbol{c}_{\boldsymbol{EI}}$ will reach nearly the same steady state, independently of the time constant.

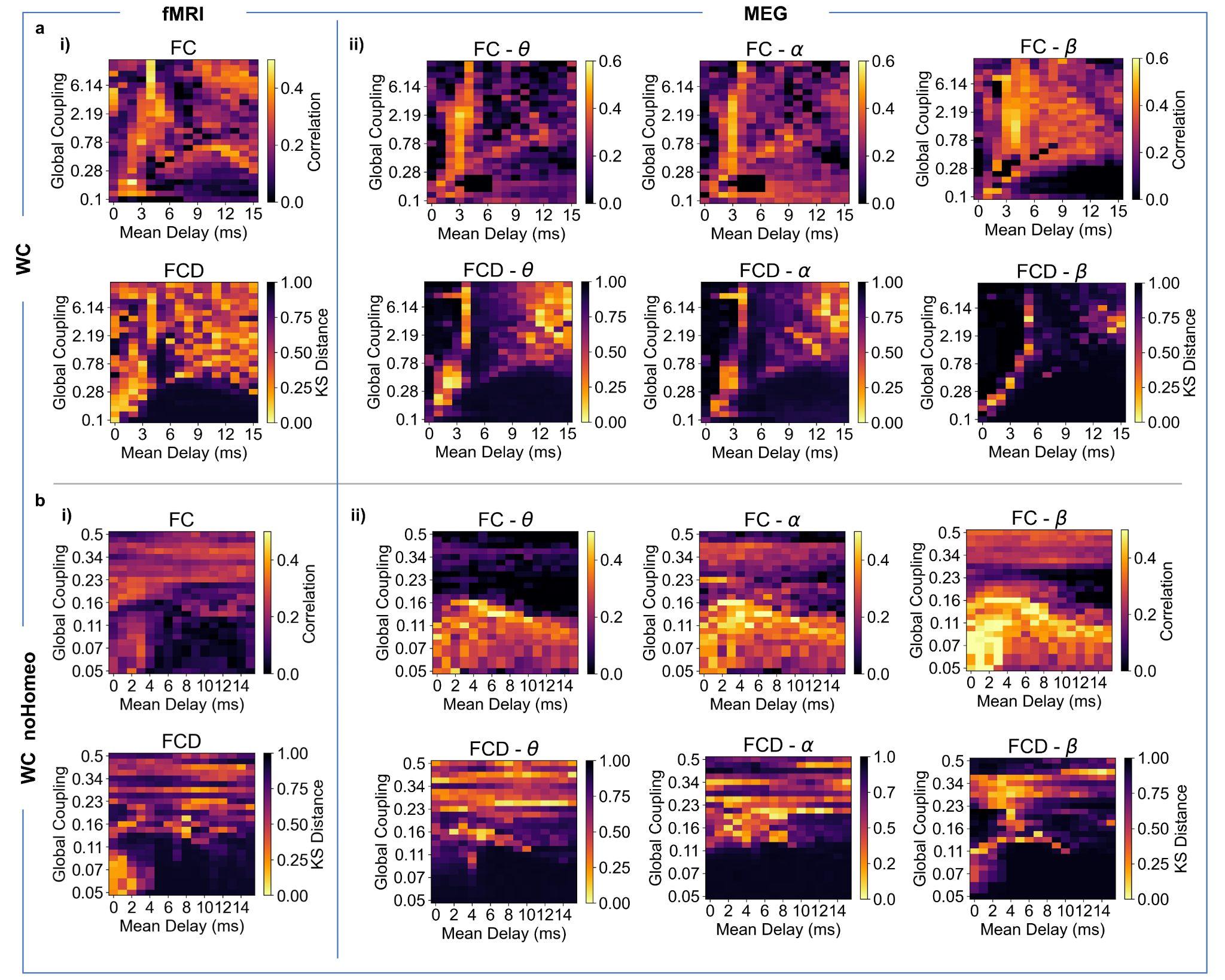

**Figure S4. WC model performance with and without plasticity. a. i)** Model performance in explaining empirical BOLD fMRI static connectivity measures: (Top) Pearson correlation between BOLD fMRI FC (averaged across 99 HCP participants) and simulated FC for each pair of parameters (Mean Delay and Global Coupling) for WC model. (Bottom) Kolmogorov-Smirnov (KS) distance between empirical BOLD fMRI FCD histograms and simulated FCD histograms for each pair of parameters for WC model. **ii)** Model performance in representing empirical MEG connectivity measures: (Top) Pearson correlation between Hilbert envelope FC of MEG (averaged across 89 HCP participants) and simulated Hilbert envelope FC for each pair of parameters, for theta [4-8 Hz] (left), alpha [8-13Hz] (middle), beta [13-30Hz] (right) for WC and SL model. (Bottom) Kolmogorov-Smirnov (KS) distance between empirical Hilbert envelope MEG FCD histograms and simulated Hilbert envelope FCD histograms for each pair of parameters, for theta [4-8 Hz] (*left*), alpha [8-13Hz] (*middle*), beta [13-30Hz] (*right*) for WC model. **b.** WC model performance without homeostatic plasticity. Same as above. In this context, the range of global couplings chosen differs because, for values of coupling higher than 0.5, most of the nodes in the network enter a saturated regime where the simulations are not physiologically plausible.

#### Target firing rate

We ran simulations also for different values of the target firing rate to investigate the importance of including a local parameter.

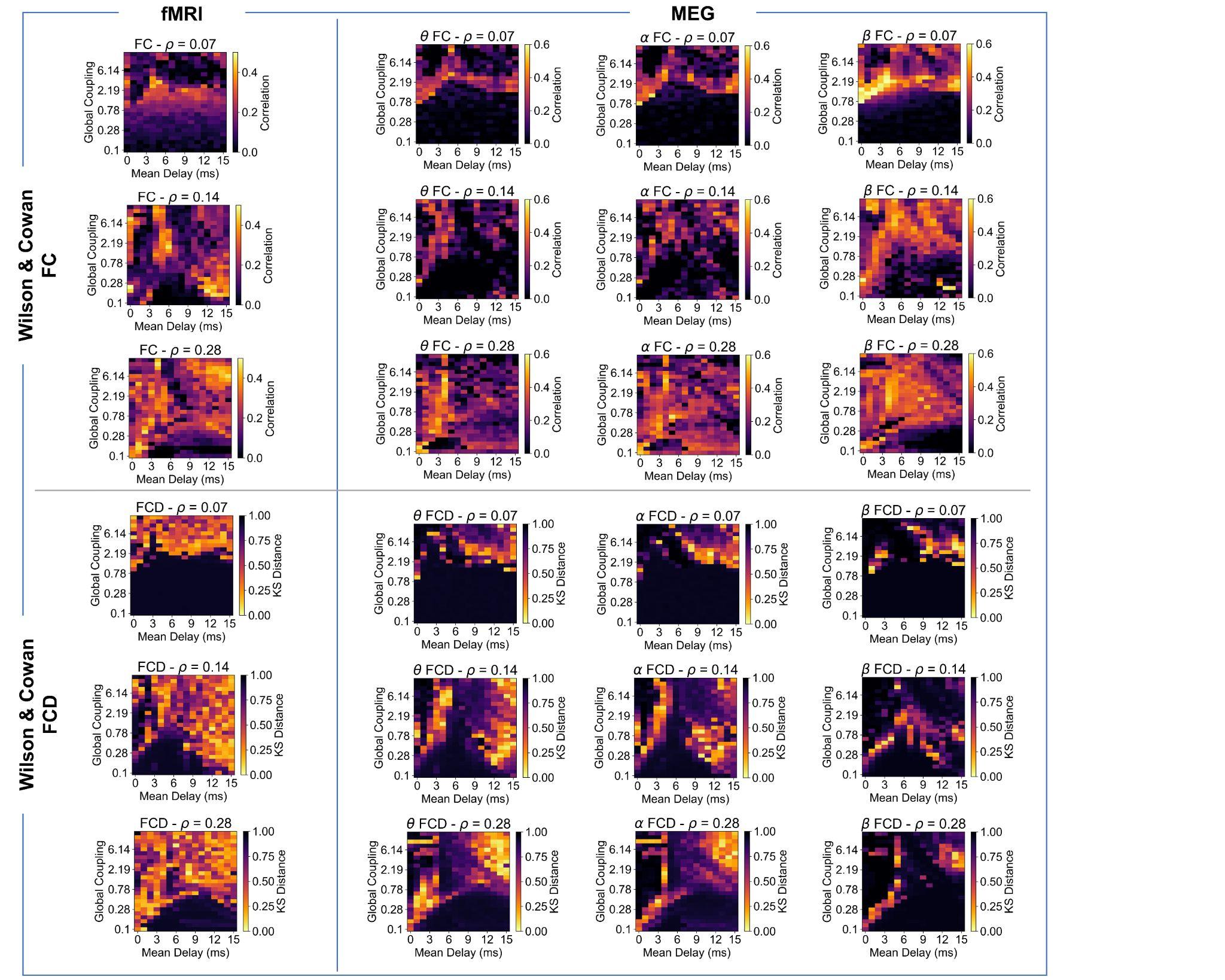

**Figure S5. Wilson and Cowan model performance for different values of the target firing rate.** Model performance in explaining empirical BOLD fMRI and MEG static and dynamic functional connectivity measures for three different values of the target firing rate $\varrho$, that is $\varrho=0.07$, $\varrho=0.14$, $\varrho=0.28$.

#### Model optimisation: evaluating the stability of the local inhibitory weights

We record $\boldsymbol{c}_{\boldsymbol{EI}}$ weights every 10s, enough to capture their slow dynamics. We then monitor the evolution of $\boldsymbol{c}_{\boldsymbol{EI}}$and allow simulations to run for either 500 minutes of simulation time or until local weights have converged to a steady state for all network nodes, evaluated through the condition described in Figure S6.

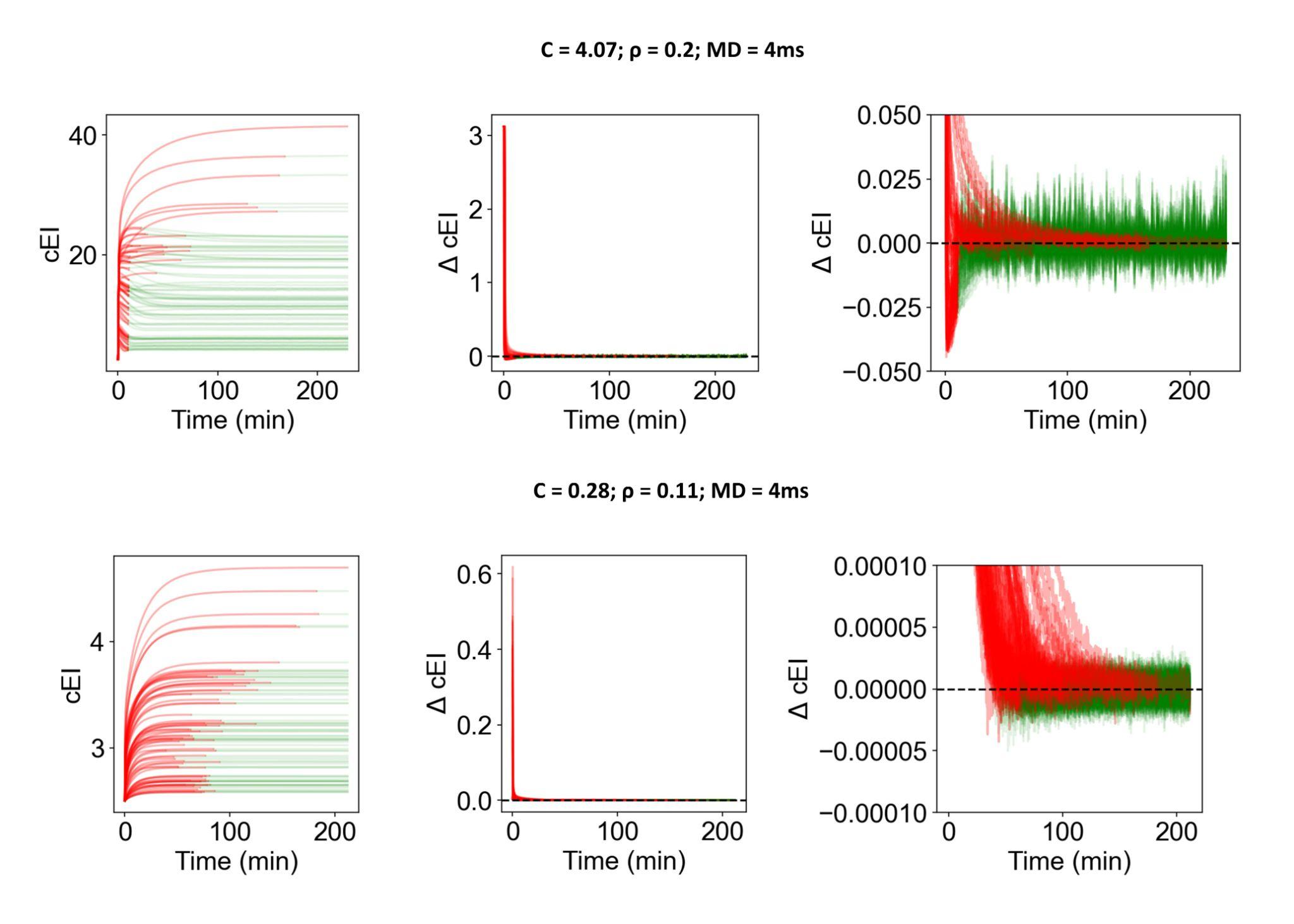

**Figure S6. Steady state test condition for** $c_{EI}$**.** Every 10 seconds, a vector keeping a down-sampled version of $c_{EI}$ in the last 10 minutes is created for every node as follows: $c_{EI,vec}(t)=[c_{EI}$(t-$T_{window}), c_{EI}(t-T_{window}+10s), ... , c_{EI}$(t-10s), $c_{EI}$(t)], where $T_{window}=600s.$ Then, The steady-state condition is fulfilled as long as the condition ${|mean(dc}_{EI,vec})|<\frac{std({dc}_{EI,vec})}{\sqrt{N}}$ remains true. When this condition is satisfied in a specific node for the first time during a simulation, we consider that node to have reached a steady state in terms of $c_{EI}$ weight. If for a specific node, the absolute mean change of $c_{EI}$ in the last 10 minutes is smaller than the standard error of the mean in the same period, the value is considered stable. Since the rate of variation of $c_{EI}$ decreases until the local firing rate gets close to the target firing rate, ${|mean(dc}_{EI,vec})|$ will decrease until it approaches 0. However, to account for the stochasticity of the system, we compare the mean variation with its respective standard error. Therefore, we effectively detect when the tendency of variation caused by homeostatic plasticity trying to restore EI balance is smaller than changes caused by the inherent stochasticity of the model.

### SECTION II

#### Selection of optimal points in the models’ parameter space

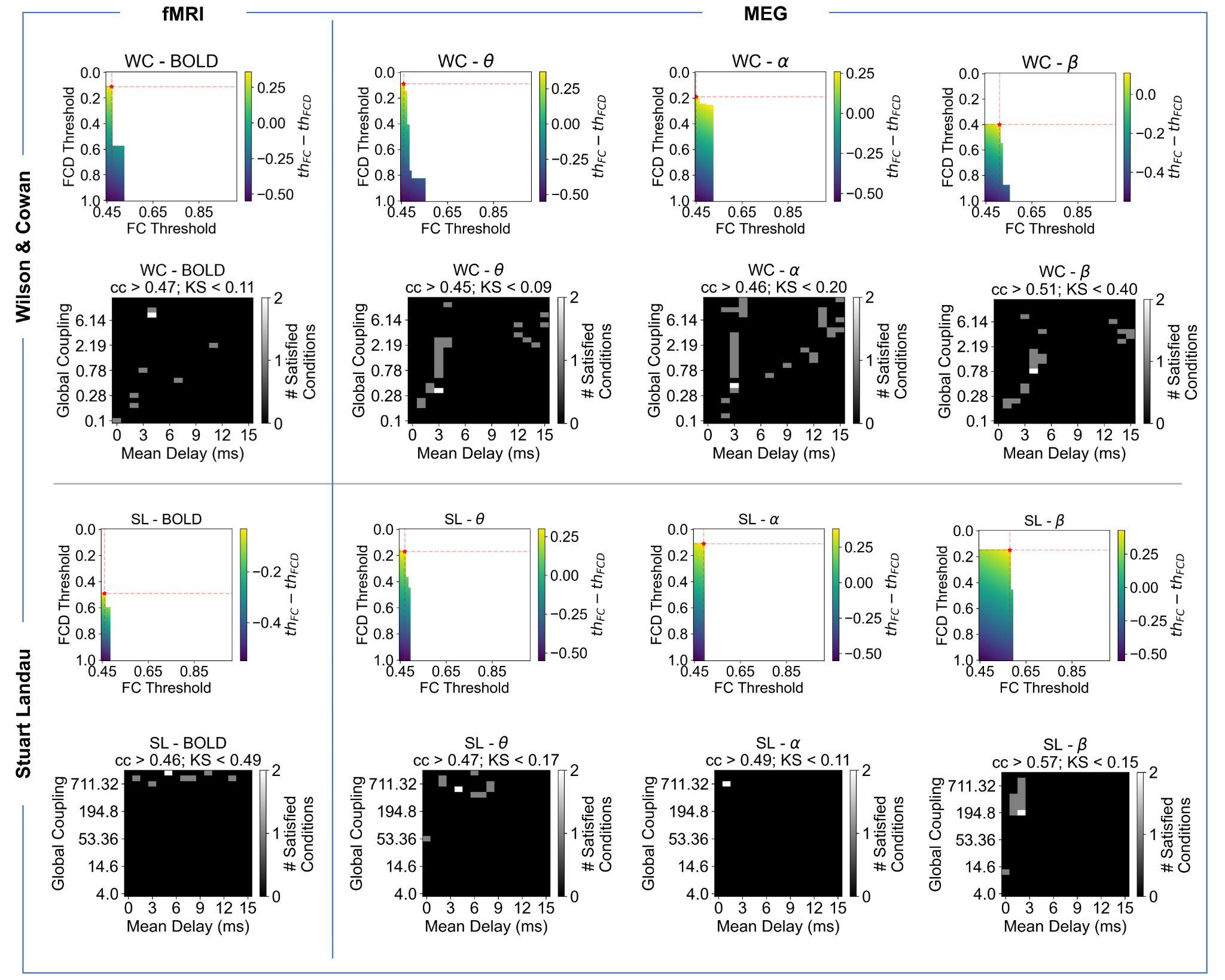

**Figure S7. Choice of optimal points across features.** Optimal point for each model and each measured modality (BOLD, MEG theta, MEG alpha and MEG beta) are chosen by iterating over a range of thresholds for FC correlation (cc ≥ ${th}_{FC}$) and FCD KS-distance (KS ≤ ${th}_{FCD}$). We then identified the maximum value of (${th}_{FC}- {th}_{FCD}$) for which both conditions can be satisfied by at least one point in the parameter space, while maximising the FC correlation and minimising the KS-distance between FCD distributions. Here, regarding FC, we impose a minimum of 0.45 since we are primarily focusing on the representation of relevant FC patterns. Red stars in the top plots indicate the chosen thresholds that maximise (${th}_{FC}- {th}_{FCD}$) and white squares in the bottom plots represent the points in the parameter space that satisfy both conditions.

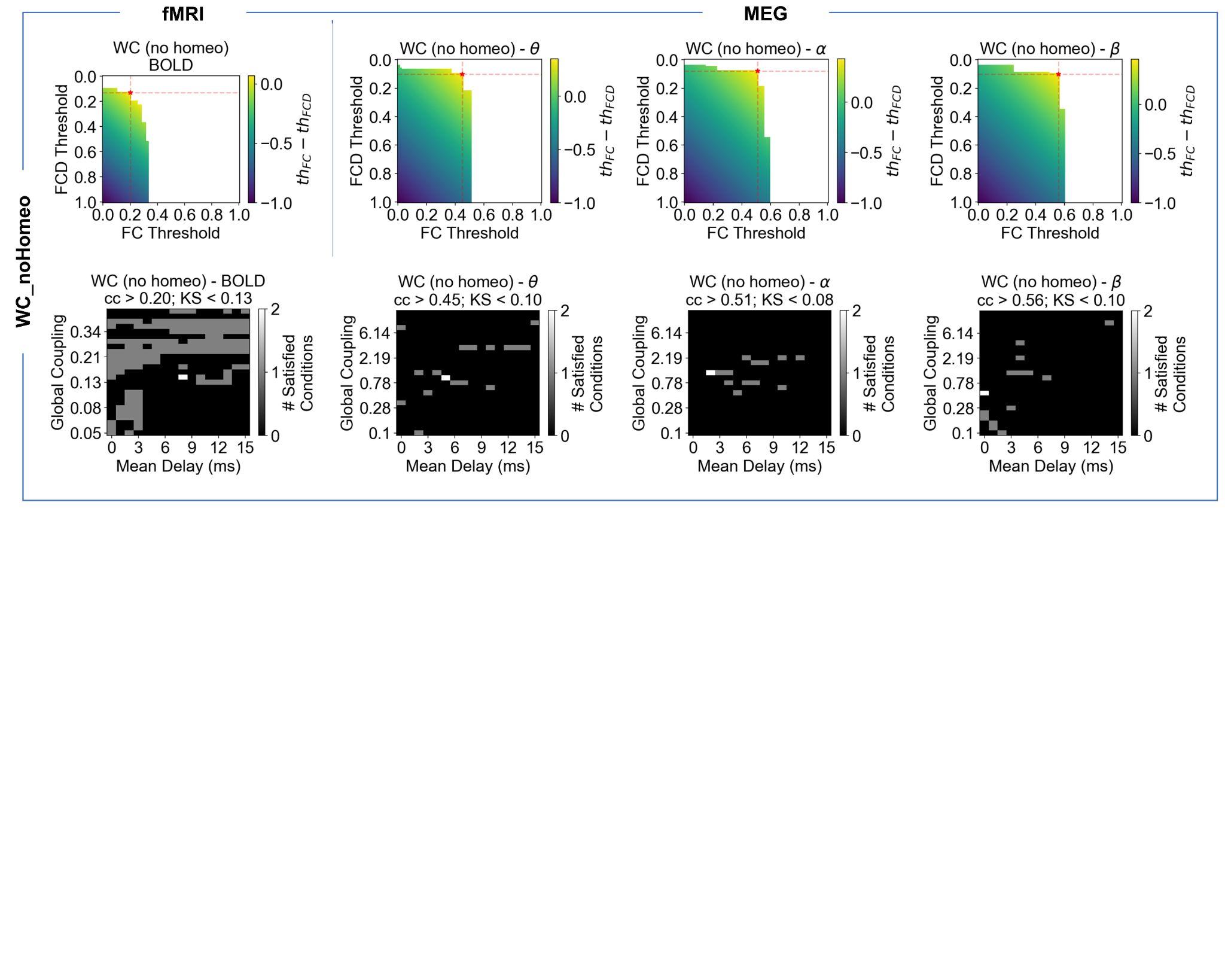

**Figure S8. Choice of optimal points for WC in the no plasticity scenario, across features.** The method used to choose the optimal points (red stars) is detailed in the Methods section and Figure S7.

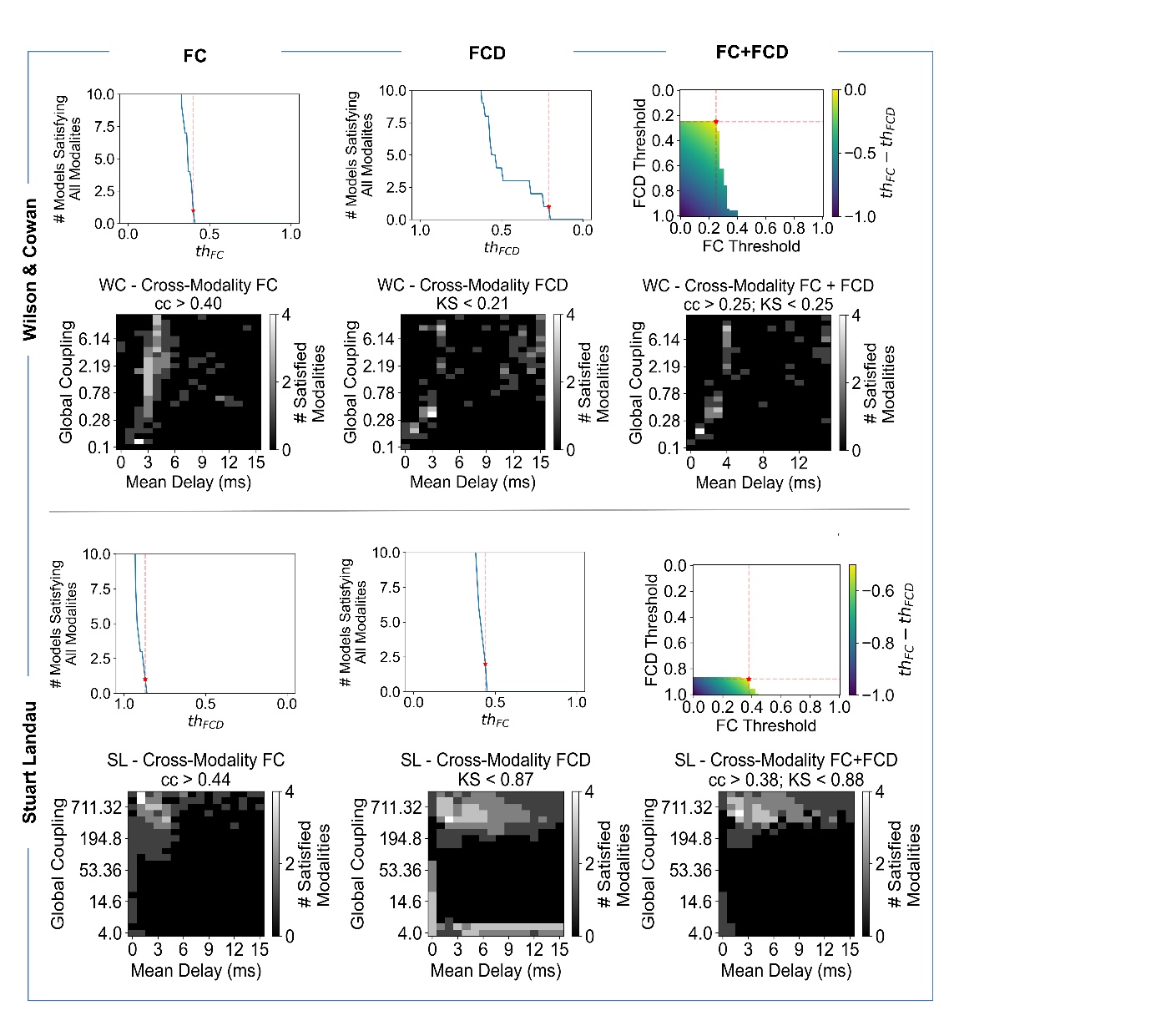

**Figure S9. Cross-modality threshold optimization (white stars in the main document – Figures 2-3).**

Optimal point for each model and each measured modality (BOLD, MEG theta, MEG alpha and MEG beta) are chosen by iterating over a range of thresholds for FC correlation (cc ≥ ${th}_{FC}$) and FCD KS-distance (KS ≤ ${th}_{FCD}$). We then identified the maximum value of (${th}_{FC}- {th}_{FCD}$) for which conditions can be satisfied across modalities by at least one point in the parameter space. White squares in the plots in the right represent the points in the parameter space that satisfy the respective conditions across all signal modalities.

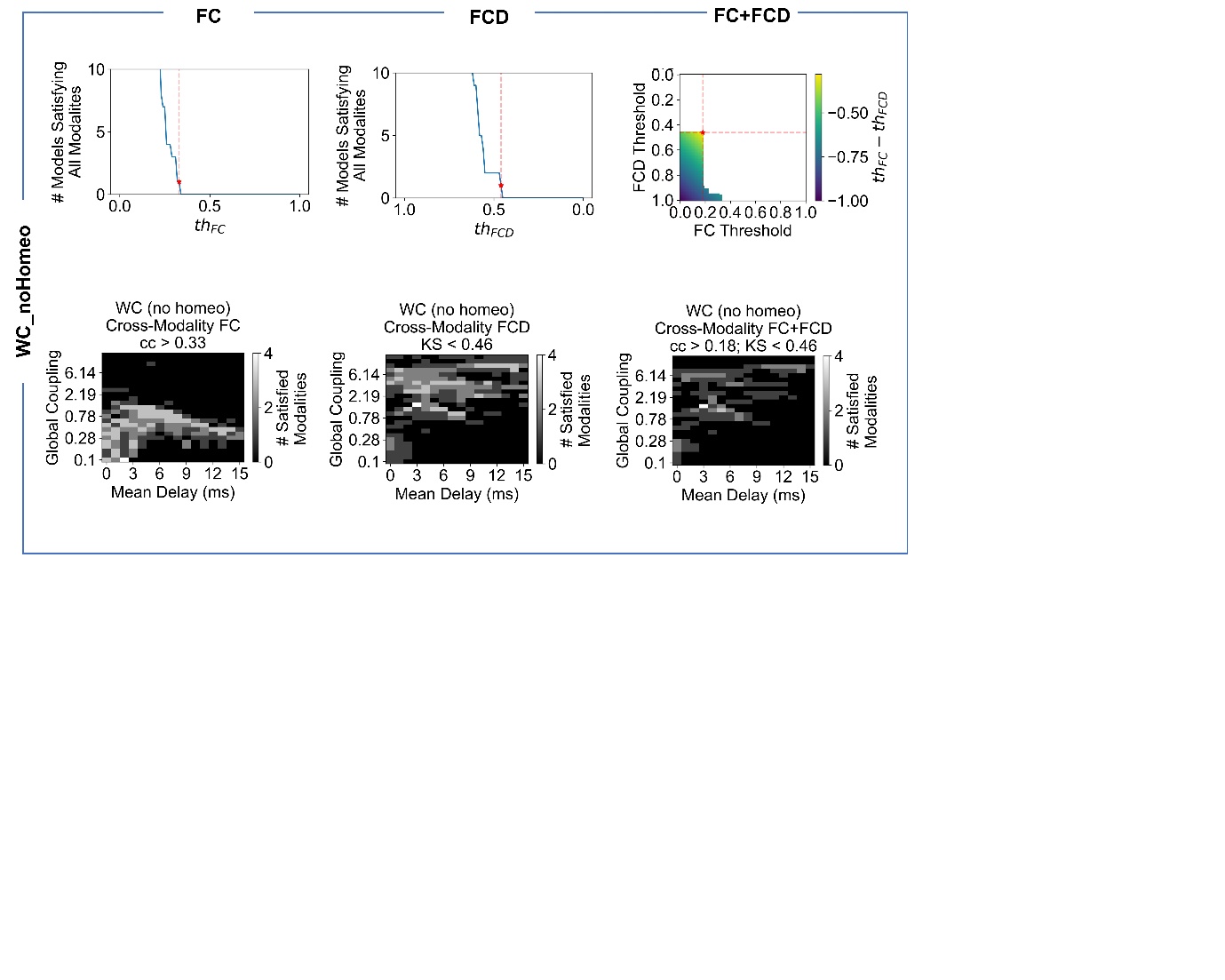

**Figure S10. Cross-modality threshold optimization in no plasticity scenario.** The algorithm to choose the optimal points for the no-plasticity scenario is the same as the one described in the Methods section of the main document and Figure S9.

#### Performance values optimised to represent FC or FCD

| **Best fit (Pearson correlation) with delays, optimised for FC only** | | | | | **Best fit (KS distance) with delays, optimised for FCD only** | | | |
| --- | --- | --- | --- | --- | --- | --- | --- | --- |
|  | **fMRI** | **Theta** | **Alpha** | **Beta** | **fMRI** | **Theta** | **Alpha** | **Beta** |
| **WC** | 0.529 | 0.550 | 0.524 | 0.555 | 0.086 | 0.035 | 0.045 | 0.061 |
| **SL** | 0.484 | 0.494 | 0.494 | 0.593 | 0.489 | 0.114 | 0.104 | 0.149 |

**Table S1. Performance values, individually optimised for FC and FCD features, accounting for delays.**

| **Best fit (Pearson correlation) without delays, optimised for FC only** | | | | | **Best fit (KS distance) without delays, optimised for FCD only** | | | |
| --- | --- | --- | --- | --- | --- | --- | --- | --- |
|  | **fMRI** | **Theta** | **Alpha** | **Beta** | **fMRI** | **Theta** | **Alpha** | **Beta** |
| **WC** | 0.436 | 0.331 | 0.300 | 0.337 | 0.082 | 0.579 | 0.712 | 0.458 |
| **SL** | 0.458 | 0.337 | 0.383 | 0.496 | 0.584 | 0.074 | 0.147 | 0.149 |

**Table S2. Performance values, individually optimised for FC and FCD features, disregarding delays.**

#### Metastable Oscillatory Modes

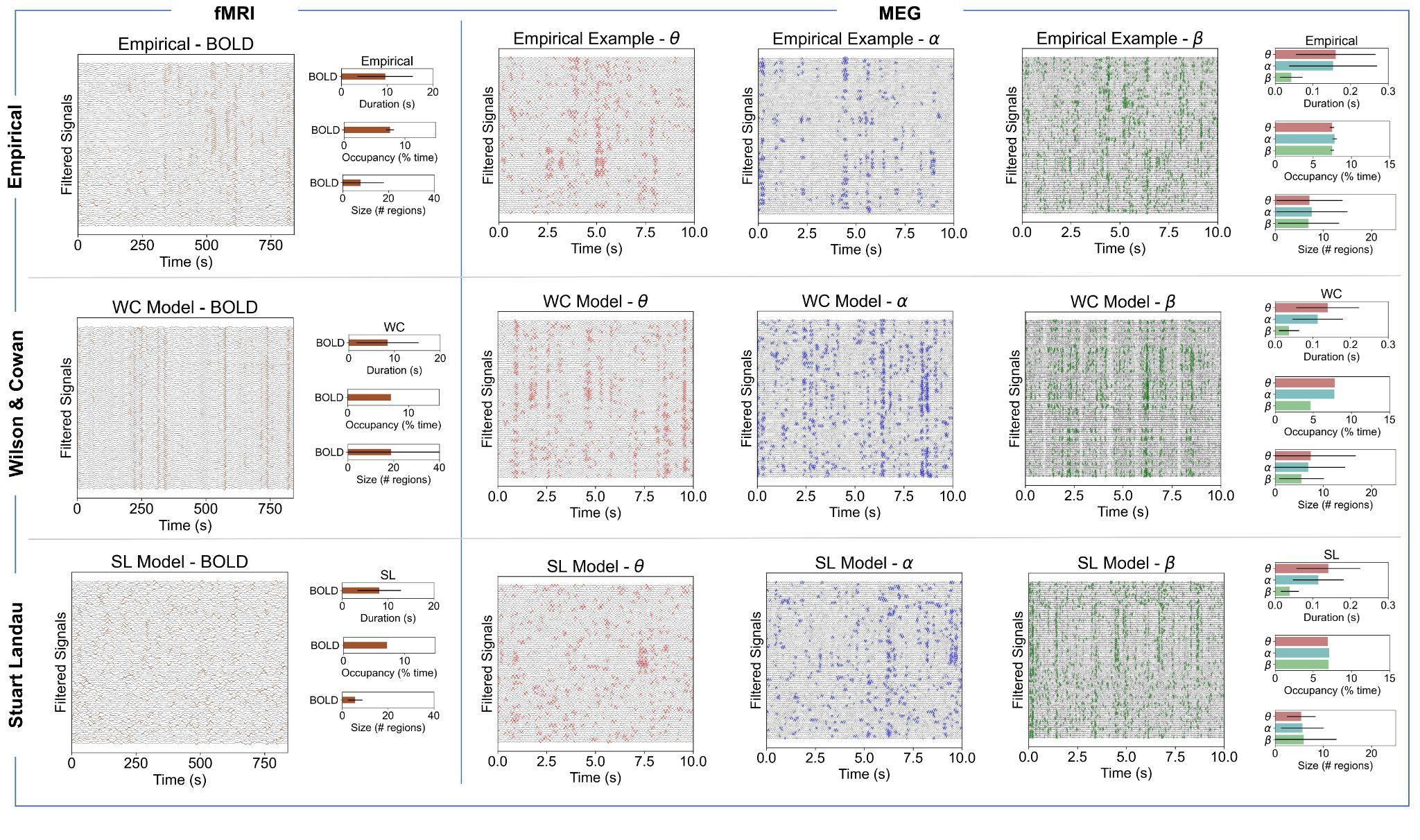

**Figure S11. Different threshold validation for MOMs detection.** Examples of empirical and simulated signals in 78 anatomically-defined regions plotted over 800s minutes for BOLD fMRI and 10 seconds for MEG. The coloured parts show the points in time when the signal power exceeds a certain threshold. For each modality, the threshold is defined as the *2.3* standard deviation of the amplitude of the signal itself. The transient itinerance is characterised in terms of duration, occupancy and size**.**

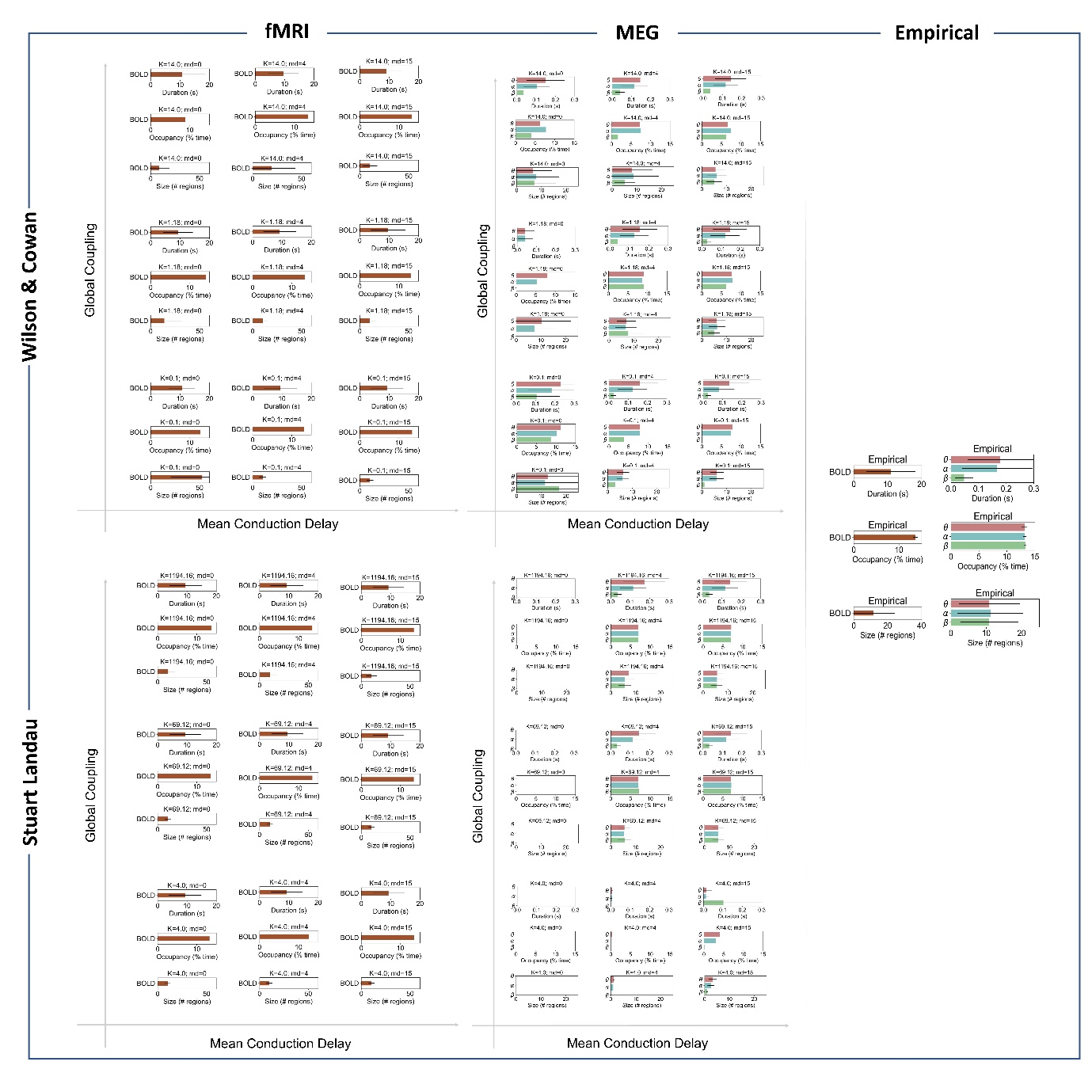

**Figure S12. Characterization of metastable oscillatory modes for different sets of models’ parameters.**

### SECTION III

#### Structural Connectivity

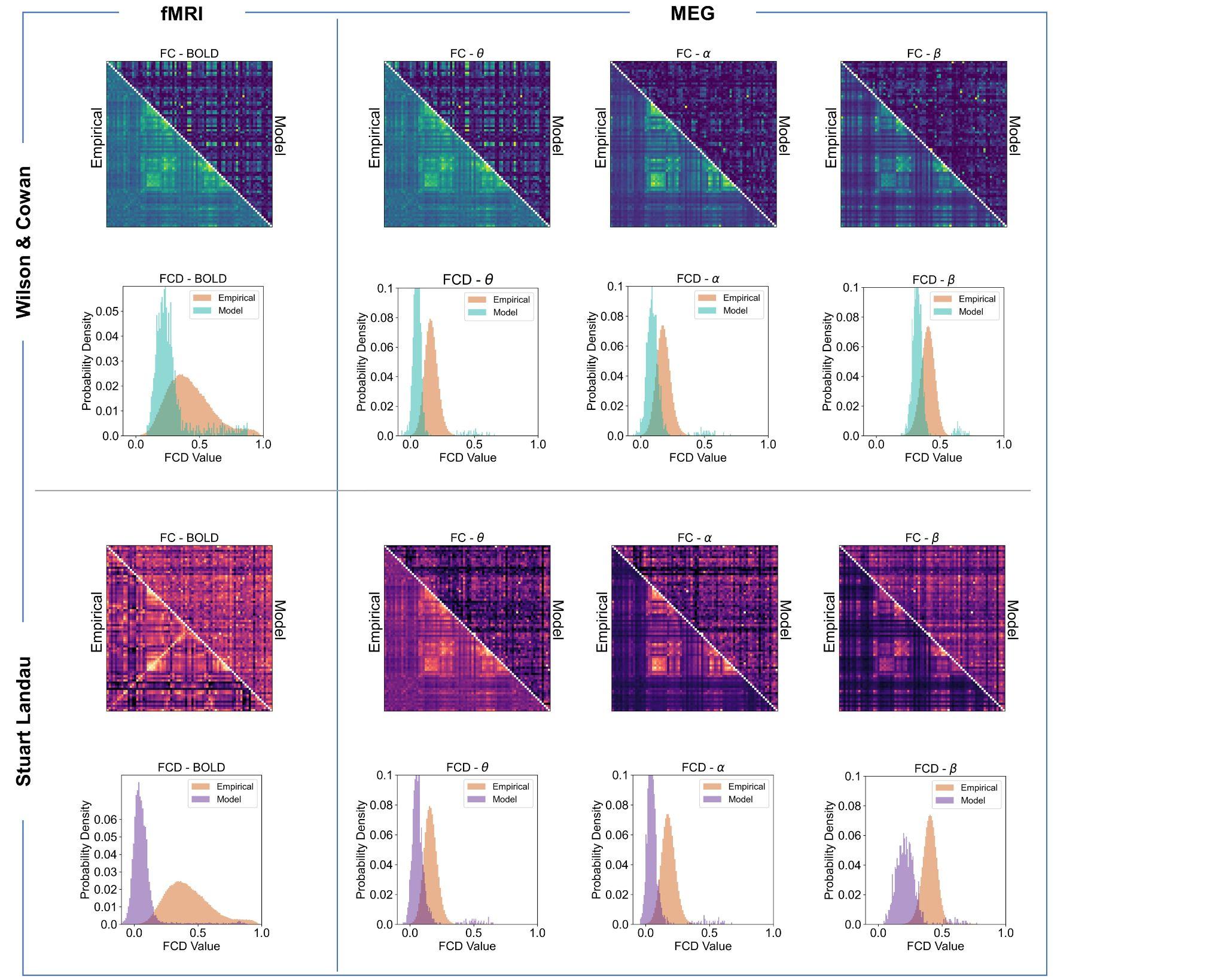

**Figure S13. WC and SL model performance giving a shuffled SC as input.** Empirical and simulated fMRI BOLD and MEG FC and FCD distributions. To prove the predictive validity of the connectome (without the right structure there is no emergence of functionally relevant patterns) we shuffled the input structural connectivity and plotted the results only for selected optimal points (white stars in Figures 2 and 3 of the main document).

### SECTION IV

#### MEG Analysis

##### Amplitude Envelope Correlation

The analytical signal vector, $z=[z_{1},z_{2} , ..., z_{T}]^{T}$ , of some temporal signal, $x=[x_{1},x_{2},...,x_{T}]^{T}$ , is defined as

$$z=x+iH[x]$$

(S1)

where $i$ is the imaginary number, and $H[x]$ is the *Hilbert* transform of the original signal. In words, the Hilbert transform provides a means of turning some real signal, x, into an analytical signal (comprising of real and imaginary parts), z, where the complex part of the analytical signal is given by the Hilbert transform of the real part of the signal. The “Hilbert Envelope”, $e=[e_{1},e_{2} , ..., e_{T}]^{T}$, of the source time series is then given by taking the absolute value of the complex output of the Hilbert transform.

If the Hilbert envelopes for brain regions A and B both have a mean of zero, the correlation between those regions is given simply by

$r_{A,B}=e_{A}e_{B}=e_{A}{e^{T}}_{B}(\sqrt{e_{A}{e^{T}}_{A}e_{B}{e^{T}}_{B}})^{-1}$,

(S2)

where $e_{A}$ and $e_{B}$ are the envelopes of brain regions A and B, respectively. Repeated application of this formula between envelopes distributed across the entire brain allows us to construct functional connectivity matrices of whole brain activity, $C_{E}\in R^{regions\times regions}$.

##### The M/EEG inverse problem and source leakage

The ill-posed nature of the inverse problem limits the spatial and temporal reconstruction accuracy of M/EEG signals, where extracranial measurements are used to approximate the activity at (typically) thousands of brain locations in a linear fashion.

In order to derive whole brain networks with M/EEG, one must address the ill-posed inverse problem. This is typically done before deriving FC metrics, although some generative FC frameworks effectively solve the inverse problem simultaneously alongside inferring FC, i.e. in Dynamic Causal Modelling (DCM) approaches (Friston et al., 2012).

Moreover, brain activity from one set of voxels can “leak” into a neighbouring set of voxels and results in artefactual zero-lag correlations featuring in reconstructed voxel time series. This leakage represents non-genuine FC between brain areas. This artefact has to be corrected for by orthogonalizing reconstructed time series via a singular value decomposition (Colclough et al., 2015) or through other means (Brookes et al., 2011; Hipp et al., 2012).

##### MEG Source Reconstruction via Beamforming

One way of tackling the M/EEG inverse problem is via beamforming. Beamformers are a data-driven spatial filters which opt to estimate a particular signal of interest arriving at some sensor array in the presence of noise (Van Veen and Buckley, 1988). A spatial filter exploits the fact that desired and interfering signals tend to originate from different positions in space. With accurate forward modelling, this spatial separation can be exploited to separate signals of interest from noise.

The general beamformer equation assumes that neural signal at voxel $i$ can be reconstructed as a weighted linear sum of sensor level measurements,

$$\hat{x_{i}(t)}={w_{i}}^{T}y(t),$$

(S3)

where $w_{i}\in R^{1\times channels}$ are the beamformer weights optimised for voxel $i$.

The linearly constrained minimum variance (LCMV) beamformer opts to minimise the overall power of the source reconstructed timeseries, with the constraint that a signal with unit amplitude originating from that voxel provides a unit response, i.e. ${w^{T}}_{i}h_{i}=1$, where $h_{i}$ is a column of the lead field matrix which describes the magnetic field that we would expect to measure at the sensor level, given a unit source placed at a specified location and orientation. The solution to these constraints is

$$w_{i}=\frac{{C^{-1}}_{Y}h_{i}}{{h^{T}}_{i}{C^{-1}}_{Y}h_{i}}$$

(S4)

where $C_{Y}\in R^{channels\times channels}$ is the empirical sensor level covariance matrix.
